# Transdiagnostic Profiles of BOLD Signal Variability in Autism and Schizophrenia Spectrum Disorders: Associations with Cognition and Functioning

**DOI:** 10.1101/2025.06.30.662421

**Authors:** Maria T. Secara, Zara Khan, Ayesha Rashidi, Lindsay D. Oliver, Ju-Chi Yu, George Foussias, Erin W. Dickie, Peter Szatmari, Pushpal Desarkar, Meng-Chuan Lai, Giulia Baracchini, Anil K. Malhotra, Robert W. Buchanan, Aristotle N. Voineskos, Stephanie H. Ameis, Colin Hawco

## Abstract

**Background:** Autism spectrum disorder (autism) and schizophrenia spectrum disorders (schizophrenia) exhibit overlapping social and neurocognitive impairment and considerable neurobiological heterogeneity. Blood-oxygen-level-dependent (BOLD) signal variability captures the brain’s moment-to-moment fluctuations, offering a dynamic marker of neural flexibility that is sensitive to cognitive capacity. This study aimed to examine intra-regional BOLD signal variability during rest and task across schizophrenia, autism, and typically developing controls (TDC) to explore transdiagnostic patterns of brain signal variability and their relationship with cognitive and functional outcomes.

**Methods:** Intra-regional BOLD variability, measured by mean squared successive difference (MSSD), was obtained from resting-state and Empathic Accuracy task fMRI in 176 SSD, 89 autism, and 149 TDC participants. ANCOVAs, controlling for age, sex, and motion, assessed group differences in regional and network-level BOLD variability and dimensional associations with social cognition, neurocognition, social functioning, and symptom severity.

**Results:** Both autism and schizophrenia exhibited lower BOLD signal variability than TDC across rest and task, with reduced variability observed in somatomotor, visual, and auditory networks (pFDR < 0.01). Greater network variability was positively associated with better social cognitive, neurocognitive, and functional scores across the sample. Resting-state variability showed stronger group-based differences and cognitive associations than task-based variability.

**Conclusions:** BOLD signal variability is positively associated with social cognition, neurocognition, and social functioning across groups, suggesting that variability impacts cognitive efficiency and behaviour. Reduced variability in autism and schizophrenia may indicate similar patterns of neural rigidity among these related conditions, positioning BOLD variability as a potential biomarker for neural flexibility and a valuable target for future transdiagnostic clinical interventions.

## 1. Introduction

Autism spectrum disorder (hereafter autism) and schizophrenia spectrum disorders (hereafter schizophrenia) are highly heritable, heterogeneous, and complex conditions. Despite being clinically distinct, both feature overlapping cognitive characteristics that contribute to poor functional outcomes and disability [1–4]. Individuals diagnosed with either condition often exhibit impairments in social cognitive [5–7] and neurocognitive domains compared to individuals without either diagnosis [8–10], with some evidence indicating that social cognitive deficits mediate the relationship between neurocognition and real-world functioning [11].

Functional neuroimaging studies of social cognition in autism and schizophrenia reveal both overlapping and distinct patterns of brain activation [7, 12–15]. However, findings are often inconsistent, potentially due to diagnostic heterogeneity and variability in sample characteristics and task demands, which may dilute or amplify observed differences [16].

The majority of imaging studies examining brain correlates of social cognition in schizophrenia and autism have examined ‘magnitude’ metrics, such as functional connectivity or task activity [17, 18]. In contrast, an fMRI metric known as timeseries variability of the blood-oxygen-level dependent (BOLD) signal provides a unique opportunity to examine the temporal unfolding of brain activity that was once deemed as noise, but is now recognized as essential to brain function [19–23]. A common measure of BOLD timeseries variability is MSSD (mean squared successive difference) [24]. Derived as the first-order derivative of the timeseries, MSSD captures moment-to-moment changes in the BOLD signal and it has been shown to be sensitive to age [25, 26], clinical status [27–31] and to reflect underlying multiscale properties of brain organisation [32]. Importantly, recent findings suggest that a brain region’s BOLD signal variability is closely related to its ‘functional embeddedness’, with greater variability observed in regions that are more strongly integrated with other brain regions [20].

Lower MSSD, indicative of reduced timeseries variability, has been related to increasing age [25, 31], and to symptom severity in neurodevelopmental conditions (e.g., attention-deficit/hyperactivity disorder and autism) [27, 28, 31]. In schizophrenia, reduced MSSD has been observed in the thalamus, sensorimotor, and visual networks, with lower dorsolateral prefrontal cortex MSSD associated with greater negative symptom severity [29]. Lower MSSD in the default-mode and salience networks has also been associated with apathy in schizophrenia, and has been linked to neural rigidity [30]. Despite these insights, most MSSD studies in clinical populations have focused on resting-state data, with task-based investigations remaining scarce [33]. Including task-based MSSD complements resting-state findings and may show greater sensitivity in detecting brain-behavior associations during cognitive engagement [34].

Building on our previous work demonstrating that social task and resting-state fMRI capture distinct, complementary aspects of individual variability in functional connectivity [35], the current study examined intra-regional BOLD variability during both task and rest across schizophrenia, autism, and TDC groups. Specifically, the objectives of this paper were to: (i) examine whether distinct group-wise patterns of intra-regional BOLD variability emerge during fMRI social cognitive task and resting-state scans, and (ii) assess dimensional brain-behavior associations between network-based BOLD variability and cognitive/clinical measures, including social cognition, neurocognition, social functioning, symptom severity, and medication use across schizophrenia, autism and TDC groups. We hypothesized that: (i) schizophrenia and autism groups would display lower regional and network-based BOLD variability compared to TDC [29–31] during rest and task, and that (ii) across our sample, lower BOLD variability would relate to reduced social cognitive and neurocognitive performance, and greater clinical symptom severity.

## 2. Methods and Materials

### 2.1 Participants

Participants, including a TDC cohort and individuals with a clinical diagnosis of schizophrenia spectrum disorder, were included from the National Institute of Mental Health (NIMH)-funded “Social Processes Initiative in the Neurobiology of Schizophrenia(s) (SPINS)” multicenter study, which, in the context of the Research Domain Criteria (RDoC) framework, investigates the clinical, behavioral, and neural correlates of social cognition across schizophrenia and controls [36–38]. Recruitment occurred at three sites: the Centre for Addiction and Mental Health (Toronto, Canada; 1/3 R01 MH102324), Zucker Hillside Hospital (Glen Oaks, NY; 2/3 R01 MH102313), and the Maryland Psychiatric Research Center (Baltimore, MD; 3/3 R01 MH102318) from 2014 to 2020. From the total SPINS sample with useable cognitive and MRI data (N = 428; TDC = 175; schizophrenia = 253, age range of 18-59), a subset of individuals aged 18-35 were included in the current analysis (N = 296; TDC = 120; schizophrenia = 176).

This age-restricted subsample was chosen to align more closely with the age-range of a prospectively recruited harmonized NIMH-funded partner study, known as the “Social Processes Initiative in the Neurobiology of Autism-spectrum and Schizophrenia-spectrum Disorders (SPIN-ASD)” (1R01MH114879-01A1,) which recruited 16-35 year old autistic and TDC participants at the Centre for Addiction and Mental Health (Toronto, Canada) and through local community agencies from 2018 to 2024. The SPIN-ASD study harmonized assessments to the SPINS study on clinical, cognitive, and imaging data collection, to expand the study of social cognition across autism, schizophrenia, and TDC groups. Information regarding participant inclusion/exclusion criteria are detailed in the Supplementary Methods. All participants provided written informed consent and the protocol was approved by the respective research ethics and institutional review boards. All research was conducted in accordance with the Declaration of Helsinki.

### 2.2 Clinical and Behavioral Assessments

In the schizophrenia and autism samples, psychiatric symptoms were assessed using the Brief Psychiatric Rating Scale (BPRS; [39]). An adapted version of the Scale for the Assessment of Negative Symptoms (SANS; [40] was used to measure negative symptoms in the schizophrenia sample. Social functioning was evaluated across schizophrenia, autism and TDCs with the Birchwood Social Functioning Scale (BSFS; [41], excluding the Employment item, and IQ was estimated using the Wechsler Test of Adult Reading (WTAR; [42] for the SPINS sample, and the Wechsler Abbreviated Scale of Intelligence – Second Edition (WASI-II, [43] in SPIN-ASD. Information regarding participant medication was collected, and chlorpromazine (CPZ) equivalents were calculated for the schizophrenia and autism groups [44].

All participants completed a set of validated social and neurocognitive measures harmonized across study protocols. The selection of social cognitive tasks aligned with SCOPE recommendations [45, 46] and included tasks that can be characterized as lower-level (e.g., emotion processing) and higher-level (e.g., theory of mind/mental state inference) social cognitive tasks. Out-of-scanner social cognitive measures included: the Penn Emotion Recognition Test (ER40; [47]) which assesses lower-level emotional recognition; the Reading the Mind in the Eyes Test (RMET; [48]), a task evaluating the recognition of complex emotions and mental states from pictures of faces depicting the eye region only; and the Awareness of Social Inference Test–Revised (TASIT) part 3 [49], which evaluates social perception through videos of actors engaged in day-to-day social situations, including scenarios involving lies and sarcasm that require integration of contextual and paralinguistic cues. From the Interpersonal Reactivity Index (IRI; [50], a self-report empathic concern subscale was included as a measure of emotional empathy, as opposed to cognitive empathy [50, 51].

The Empathic Accuracy (EA) task [52], completed during fMRI scanning [53, 54], involved participants watching 9 short videos (average length = 2.05 min, range 2.00 to 2.50 mins), presented in three runs. Participants provided continuous ratings of how positive or negative actors relating a biographical story were feeling (ranging from 1, very negative, to 9, very positive). The EA score is the correlation of the participant’s ratings with the actors/target’s self-reported feeling ratings [52]. See the Supplementary Materials for additional details.

Neurocognition was assessed using the Measurement and Treatment Research to Improve Cognition in Schizophrenia (MATRICS) Consensus Cognitive Battery (MCCB, [55]). The battery evaluates seven cognitive domains: processing speed, reasoning and problem solving, attention/vigilance, working memory, verbal learning, visual learning, and social cognition. A neurocognitive composite score was derived by averaging t-scores across MCCB tasks, excluding the social cognitive domain, to assess overall neurocognition across groups [3, 56, 57].

### 2.3 MRI Data Acquisition

Scanning was conducted on six 3T scanners with multichannel head coils and harmonized scanning parameters (see Supplementary Material). Participants from the SPIN-ASD study were scanned on one scanner (CMH), whereas SPINS participants were scanned on six scanners (CMH, CMP, MRC, MRP, ZHH and ZHP). The EA task was incorporated within a comprehensive multimodal MRI protocol, conducted as part of the harmonized SPINS and SPIN-ASD studies [36]. Three EA fMRI runs were acquired using an echo-planar imaging (EPI) sequence (TR = 2000 ms, TE = 30 ms, flip angle = 77°, field of view = 218 mm, in-plane resolution = 3.4 mm^2^, slice thickness = 4 mm). One seven-minute resting-state scan was acquired using an accelerated EPI sequence (TR = 2000 ms, TE = 30 ms, flip angle = 77°, field of view = 200 mm, in-plane resolution = 3.125 mm^2^, slice thickness = 4 mm) where participants were instructed to let their mind wander with their eyes closed for the duration of the scan. Anatomical T1-weighted scans were collected using a fast-gradient sequence (CMH and ZHH used a BRAVO sequence in sagittal plane with TR = 6.4/6.7 ms, TE = 2.8/3 ms, flip angle = 8°, voxel-size =0.9 mm; CMP, MRC, MRP and ZHP used a MPRAGE sequence in sagittal plane with TR =2300 ms, TE = 2.9 ms, flip angle = 9°, voxel-size = 0.9mm).

All scans underwent quality control via an in-house quality control system dashboard (https://github.com/TIGRLab/dashboard) and were reviewed by research staff using quantitative (e.g., mean framewise displacement; FD) and qualitative measures (e.g., ringing or ghosting).

Participants with excessive motion (mean FD > 0.5 mm) or EA/control task performance indicative of disengagement or comprehension issues were excluded (see Fig. S1 and Supplementary Material).

### 2.4 fMRI Preprocessing

Scans were preprocessed using fMRIPrep 20.2.7 [58], based on Nipype 1.7.0 [59]. Full details of the preprocessing pipeline are presented in the Supplementary Material. In brief, structural MRI was run through Freesurfer (6.0.1) to segment the cortex and subcortical structures and calculate the non-linear transform to MNI space. fMRI data underwent fieldmap-less distortion correction, were realigned for motion, and slice time corrected, and coregistered to the T1. fMRI data were then registered to the cortical surface in the Human Connectome CIFTI space using the Freesurfer tissue segmentation and nonlinear registration [60]. Scans were surface smoothed at 2mm. For both resting-state and EA data, the nuisance regression model included regressors for the six head motion correction parameters, mean white matter signal, mean cerebral spinal fluid signal, global brain signal, and the square, derivative, and square of the derivative for each of these regressors [61, 62]. Average timeseries were extracted from 360 cortical regions from the surface-based Multimodal Parcellation (MMP) 1.0 atlas [63, 64] and 32 subcortical regions from the Melbourne Subcortex Atlas (Scale II) [65]. For EA, only time points during the EA videos were retained. Networks were defined according to the Cole-Anticevic functional network organization [64], with subcortical regions grouped separately.

### 2.5 Intra-Regional BOLD Signal Variability Calculation

Variability calculations were completed using Python v3.10.12 in Jupyter Notebook v7.2.2. MSSD was used to calculate intra-regional BOLD signal variability given its ability to capture variability in signal amplitude between successive time points [24], independence from shifts in the mean [66], and proven reliability [67]. Each regional timeseries was mean-centered to remove baseline offsets. MSSD in each region was calculated by subtracting each time point from its successive time point, and then calculating the square root of the average of all subtractions for each timeseries (Fig. 1) [20]. Following MSSD calculation, neuroCombat in RStudio v2024.04 was used to remove the effects of different MRI scanners, preserving variance related to age, sex, and diagnosis [68, 69]. Network MSSD values were derived by averaging all regional MSSD values within the 12 Cole-Anticevic functional networks. This gave one MSSD value per network, representing BOLD signal variability at the network level. This parcellation integrates resting-state functional connectivity and multimodal neuroimaging data, ensuring anatomically and functionally meaningful delineation of cortical networks, addressing a key limitation of prior parcellations [64].

**Figure 1.**
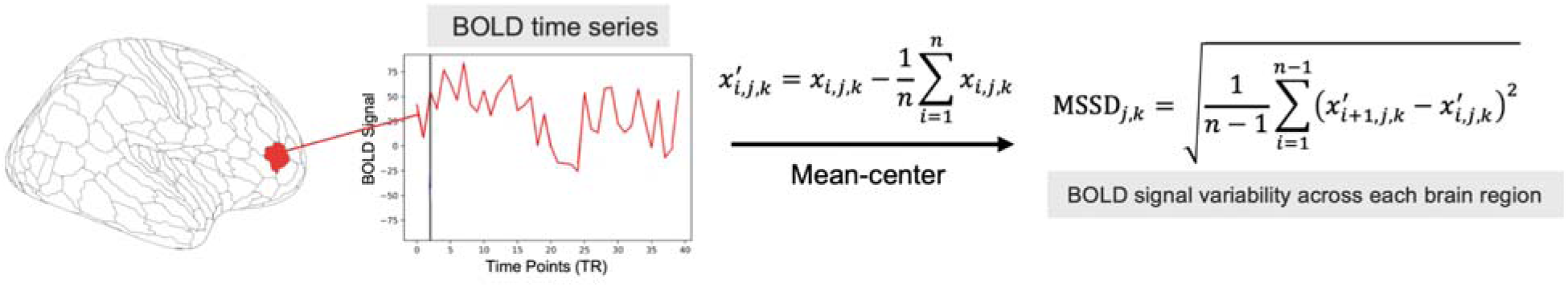
Intra-regional BOLD signal variability calculation via MSSD: Stepwise representation of timeseries variability calculation using MSSD for resting-state and EA task. Each regional timeseries was mean-centered: For each brain region *j* (where *j* = 1, 2,… 392) and participant *k,* where x_i,j,k_ is the original time series value at time point *i* for region *j* and participant *k*, and x’_i,j,k_ is the mean-centered value. Following this, a regional MSSD value was derived for all 392 brain regions on the mean-centered timeseries. Each participant received 392 MSSD values representing whole-brain timeseries variability (excluding the cerebellum), separately for resting-state and EA task.

### 2.7 Statistical Analysis

Analyses were completed in RStudio v2024.04.2 and evaluated separately for resting-state and EA task timeseries variability. Regional-and network-based MSSD values were compared between autism, schizophrenia and TDC using one-way ANCOVAs including age, sex assigned at birth, and mean FD as covariates. Multiple comparisons were corrected using false discovery rate (FDR) (q < 0.05) across 392 brain regions and 12 networks. Tukey HSD post hoc tests were conducted to identify pairwise group differences in significant ANCOVAs.

#### 2.7.1 Relationship between Network MSSD and Behavioral/Cognitive/Clinical Variables

Given the prior associations found between timeseries variability, cognition, symptom severity, and behavior [21, 25, 27–30, 66], we conducted one-way ANCOVAs across groups to examine whether network-based variability was associated with key cognitive, clinical and behavioral variables. Age, sex, and mean FD were included as covariates, and FDR correction (*q* < 0.05) was applied across all tested network–cognitive/outcome variable associations. Key cognitive variables included: social cognition (TASIT3 Sarcasm, TASIT3 Lies, ER40 total, and IRI empathic concern score), neurocognition (MCCB composite score), social functioning (BSFS total score), symptom severity (BPRS total score), and medication effects (CPZ equivalents); the relationship between CPZ equivalents and network variability was examined to assess potential medication-related contributions to group differences. Social cognition measures were selected to provide measures of both: lower-level social cognition, involving emotional recognition (ER40 total), as well as measures involving more complex social cognition tasks, such as detecting lies and sarcasm (TASIT3 Lies score, TASIT3 Sarcasm score). In particular, the TASIT3 sarcasm score was selected to assess higher-level social cognition [11]. Notably, the TASIT3 sarcasm score is specifically associated with theory of mind [70] and loads strongly onto a higher-level mentalizing factor in schizophrenia [11]. Mancuso et al further demonstrated that theory of mind is a key component of social cognition in psychosis, with poorer TASIT3 sarcasm performance in schizophrenia linked to greater positive symptoms and reduced social functioning [70]. To further explore whether these associations differed across groups, post hoc ANCOVAs including interaction terms between group and network variability (e.g., network × group) were conducted. These analyses were conducted across schizophrenia, autism, and TDC groups during both rest and task conditions.

### 2.8 Code Sharing

Code used in the analysis of this dataset has been made available (https://github.com/tsecara/BOLD_signal_variability_Autism_Schizophrenia_TDC).

## 3. Results

### 3.1 Participant Inclusion and Characteristics

Data from 176 individuals with schizophrenia, 89 with autism and 149 TDC (total N = 414) were included from the SPINS and SPASD studies (Fig. S1). Participant demographics and clinical characteristics, as well as clinical, social cognitive and neurocognitive scores, are shown in Table 1. A comprehensive breakdown of medication usage is detailed in Supplemental Table S1.

**Table 1.**
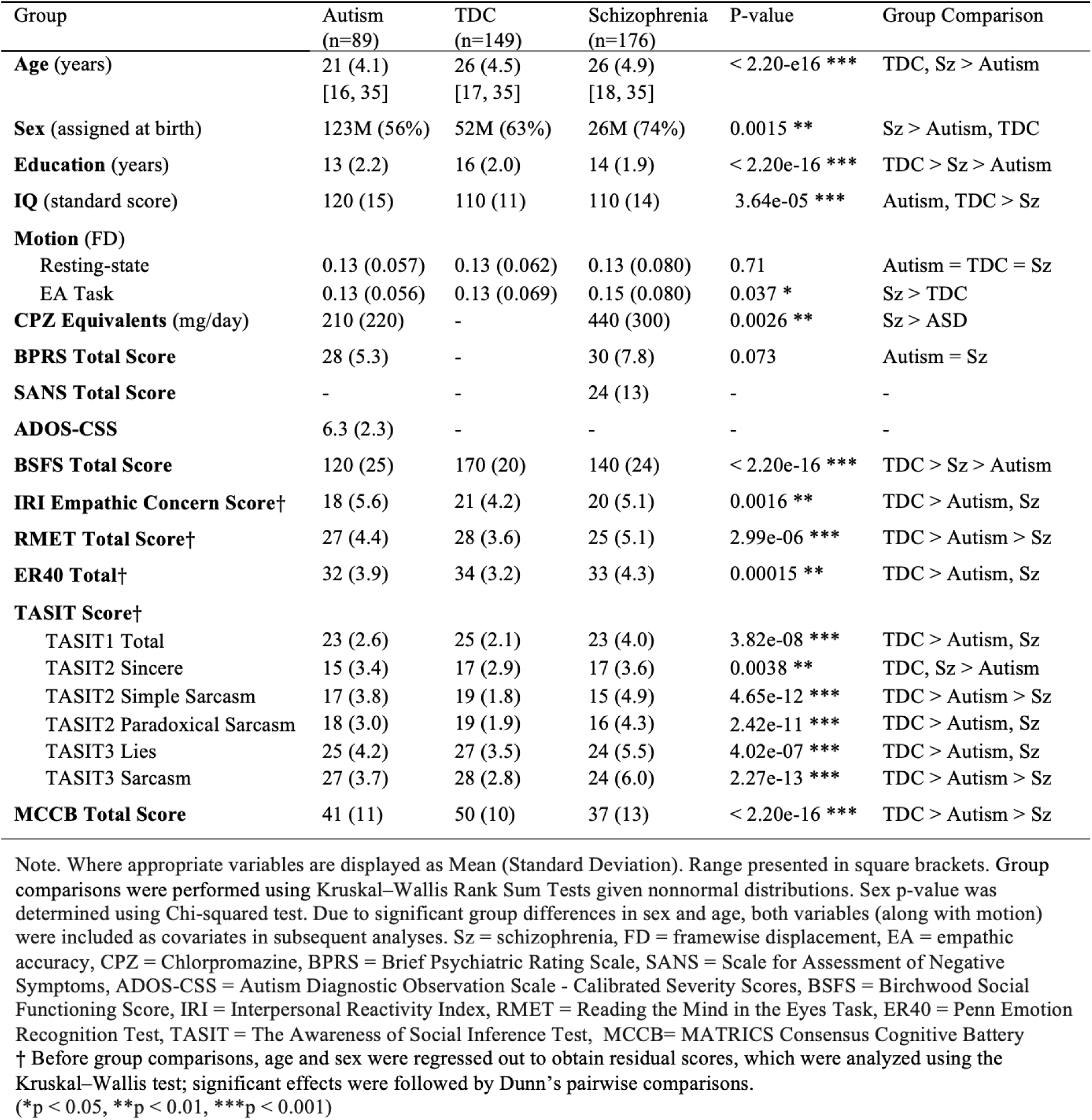
Participant Demographic and Clinical Characteristics by Diagnostic Group.

| Group | Autism<br>(n=89) | TDC<br>(n=149) | Schizophrenia<br>(n=176) | P-value | Group Comparison |
| --- | --- | --- | --- | --- | --- |
| **Age (years)** | 21 (4.1)<br>[16, 35] | 26 (4.5)<br>[17, 35] | 26 (4.9)<br>[18, 35] | < 2.20e-16 *** | TDC, Sz > Autism |
| **Sex (assigned at birth)** | 123M (56%) | 52M (63%) | 26M (74%) | 0.0015 ** | Sz > Autism, TDC |
| **Education (years)** | 13 (2.2) | 16 (2.0) | 14 (1.9) | < 2.20e-16 *** | TDC > Sz > Autism |
| **IQ (standard score)** | 120 (15) | 110 (11) | 110 (14) | 3.64e-05 *** | Autism, TDC > Sz |
| **Motion (FD)** |  |  |  |  |  |
| Resting-state | 0.13 (0.057) | 0.13 (0.062) | 0.13 (0.080) | 0.71 | Autism = TDC = Sz |
| EA Task | 0.13 (0.056) | 0.13 (0.069) | 0.15 (0.080) | 0.037 * | Sz > TDC |
| **CPZ Equivalents (mg/day)** | 210 (220) | - | 440 (300) | 0.0026 ** | Sz > ASD |
| **BPRS Total Score** | 28 (5.3) | - | 30 (7.8) | 0.073 | Autism = Sz |
| **SANS Total Score** | - | - | 24 (13) | - | - |
| **ADOS-CSS** | 6.3 (2.3) | - | - | - | - |
| **BSFS Total Score** | 120 (25) | 170 (20) | 140 (24) | < 2.20e-16 *** | TDC > Sz > Autism |
| **IRI Empathic Concern Score†** | 18 (5.6) | 21 (4.2) | 20 (5.1) | 0.0016 ** | TDC > Autism, Sz |
| **RMET Total Score†** | 27 (4.4) | 28 (3.6) | 25 (5.1) | 2.99e-06 *** | TDC > Autism > Sz |
| **ER40 Total†** | 32 (3.9) | 34 (3.2) | 33 (4.3) | 0.00015 ** | TDC > Autism, Sz |
| **TASIT Score†** |  |  |  |  |  |
| TASIT1 Total | 23 (2.6) | 25 (2.1) | 23 (4.0) | 3.82e-08 *** | TDC > Autism, Sz |
| TASIT2 Sincere | 15 (3.4) | 17 (2.9) | 17 (3.6) | 0.0038 ** | TDC, Sz > Autism |
| TASIT2 Simple Sarcasm | 17 (3.8) | 19 (1.8) | 15 (4.9) | 4.65e-12 *** | TDC > Autism > Sz |
| TASIT2 Paradoxical Sarcasm | 18 (3.0) | 19 (1.9) | 16 (4.3) | 2.42e-11 *** | TDC > Autism, Sz |
| TASIT3 Lies | 25 (4.2) | 27 (3.5) | 24 (5.5) | 4.02e-07 *** | TDC > Autism, Sz |
| TASIT3 Sarcasm | 27 (3.7) | 28 (2.8) | 24 (6.0) | 2.27e-13 *** | TDC > Autism > Sz |
| **MCCB Total Score** | 41 (11) | 50 (10) | 37 (13) | < 2.20e-16 *** | TDC > Autism > Sz |

Note. Where appropriate variables are displayed as Mean (Standard Deviation). Range presented in square brackets. Group comparisons were performed using Kruskal–Wallis Rank Sum Tests given nonnormal distributions. Sex p-value was determined using Chi-squared test. Due to significant group differences in sex and age, both variables (along with motion) were included as covariates in subsequent analyses. Sz = schizophrenia, FD = framewise displacement, EA = empathic accuracy, CPZ = Chlorpromazine, BPRS = Brief Psychiatric Rating Scale, SANS = Scale for Assessment of Negative Symptoms, ADOS-CSS = Autism Diagnostic Observation Scale - Calibrated Severity Scores, BSFS = Birchwood Social Functioning Score, IRI = Interpersonal Reactivity Index, RMET = Reading the Mind in the Eyes Task, ER40 = Penn Emotion Recognition Test, TASIT = The Awareness of Social Inference Test, MCCB= MATRICS Consensus Cognitive Battery
† Before group comparisons, age and sex were regressed out to obtain residual scores, which were analyzed using the Kruskal–Wallis test; significant effects were followed by Dunn’s pairwise comparisons.
(\*p < 0.05, \*\*p < 0.01, \*\*\*p < 0.001)

### 3.2 Regional-and Network-based Variability

Across groups, MSSD was generally greater in association regions (e.g., within the default mode network), and lower in sensory-motor regions, with similar patterns for MSSD values among cortical regions for task and rest-derived data (Fig. 2A and 2C), in line with previously reported findings [29, 67, 71].

**Figure 2.**
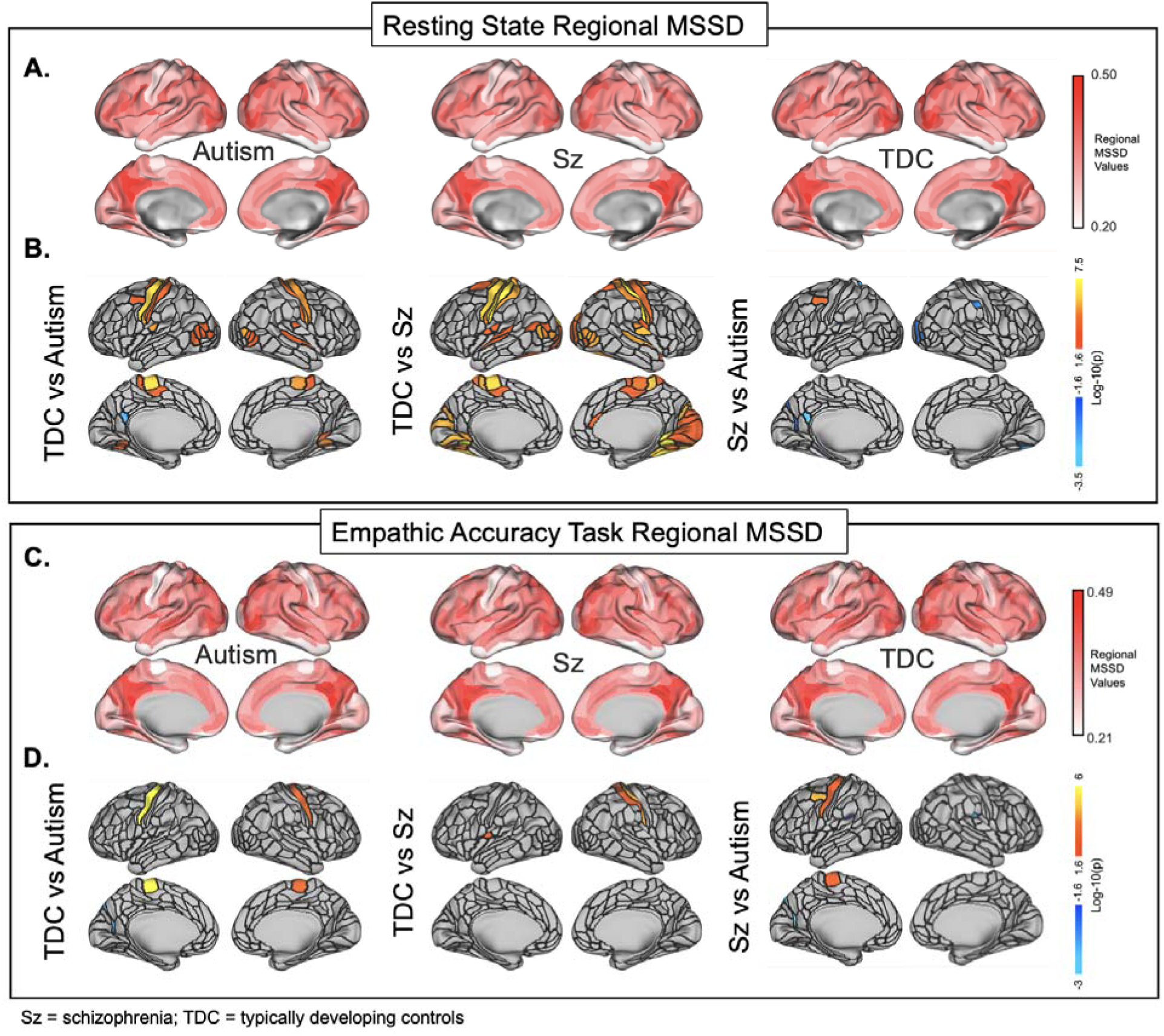
Regional Variability via MSSD: A&C) Regional representation of brain variability across autism, schizophrenia and TDC during resting-state and EA task. Darker red regions depict greater regional variability. **B&D)** Group comparison of regional variability across groups during resting-state and EA task. From left to right: orange/red regions depict areas of the brain where TDC show increased variability compared to autism; orange/red regions depict areas of the brain where TDC show increased variability compared to schizophrenia; orange/red regions depict areas of the brain where schizophrenia show increased variability compared to autism and blue regions depict where autism group show increased variability compared to schizophrenia.

#### 3.2.1 Group-Wise Differences in Resting-State Regional Timeseries Variability

Group-level findings indicated that TDC groups exhibited significantly higher MSSD across multiple regions, prominently involving sensory-motor regions, compared to both schizophrenia and autism groups, after FDR correction (Fig. 2B). TDC displayed greater MSSD (i.e., higher variability) compared to schizophrenia across 25% of all regions examined (96/392 regions; 5 subcortical and 91 cortical). TDC displayed greater MSSD compared to the autism group in 12% of regions (46/392 regions; 7 subcortical and 39 cortical). The autism group had greater MSSD in one cortical region (Area ventral 23 a+b L in the posterior cingulate) when compared to TDC. When comparing schizophrenia versus autism groups, 5% of regions were significantly different (18/392 regions; 3 subcortical and 15 cortical). Group differences between schizophrenia and autism were primarily driven by greater timeseries variability in the autism group, observed in 14 out of 18 regions (78%) that showed significant differences. These 14 regions included: subcortical areas (anterior caudate, globus pallidus), visual regions (V4, V8, LO1, VVC), parietal regions (7AL, 7Am), and an auditory region (LBelt). Greater MSSD was found in the schizophrenia group compared to autism across 4 regions, including the anterior caudate (left and right; aCAU.lh, aCAU.rh), left FST (Fusiform Sulcus Temporalis), and left area 55b, a region in the posterior inferior frontal gyrus linked to language and cognitive control.

#### 3.2.2 Group-Wise Differences in EA task Derived Regional Timeseries Variability

Consistent with resting-state findings, TDC exhibited higher MSSD than schizophrenia and autism for EA task-derived timeseries data. However, regional group differences were observed in fewer areas compared to resting-state (Fig. 2D). TDC displayed higher MSSD compared to schizophrenia in 1% of regions (4/392 regions; 1 subcortical and 3 cortical). TDC displayed higher MSSD compared to the autism group in 4% of regions (14/392 regions; 3 subcortical and 11 cortical). The autism group featured higher MSSD in one cortical region (Dorsal translation visual area L in the posterior cingulate) when compared to TDC. For schizophrenia-autism group comparisons, 3% of regions were significantly different (12/393 regions; 3 subcortical and 9 cortical). Increased regional MSSD was found in 7/12 regions in schizophrenia versus autism, including regions encompassing the nucleus accumbens (bilaterally), anterior cingulate cortex (bilaterally), primary motor cortex, and the premotor area. Increased regional MSSD was found in 5/12 regions in autism versus schizophrenia, including regions encompassing the inferior parietal cortex (bilaterally), dorsal visual stream, ventral temporal gyrus, and the insular cortex.

#### 3.2.3 Group-Wise Differences in Network Timeseries Variability During Rest and EA Task

Significant group differences were detected in brain networks during rest (Fig. 3A), in unimodal networks including: the visual 1, visual 2, somatomotor and auditory networks, as well as in the cingulo-opercular network (outlined in Table S3). TDC displayed greater MSSD compared to schizophrenia in the subcortical, visual 1, visual 2, somatomotor, cingulo-opercular, and auditory networks. TDC displayed greater MSSD compared to the autism group in the subcortical, visual 2, somatomotor, and auditory networks. In addition to differences found between TDC and the two clinical groups, MSSD was greater in the autism versus schizophrenia group within the visual 1 network.

**Figure 3.**
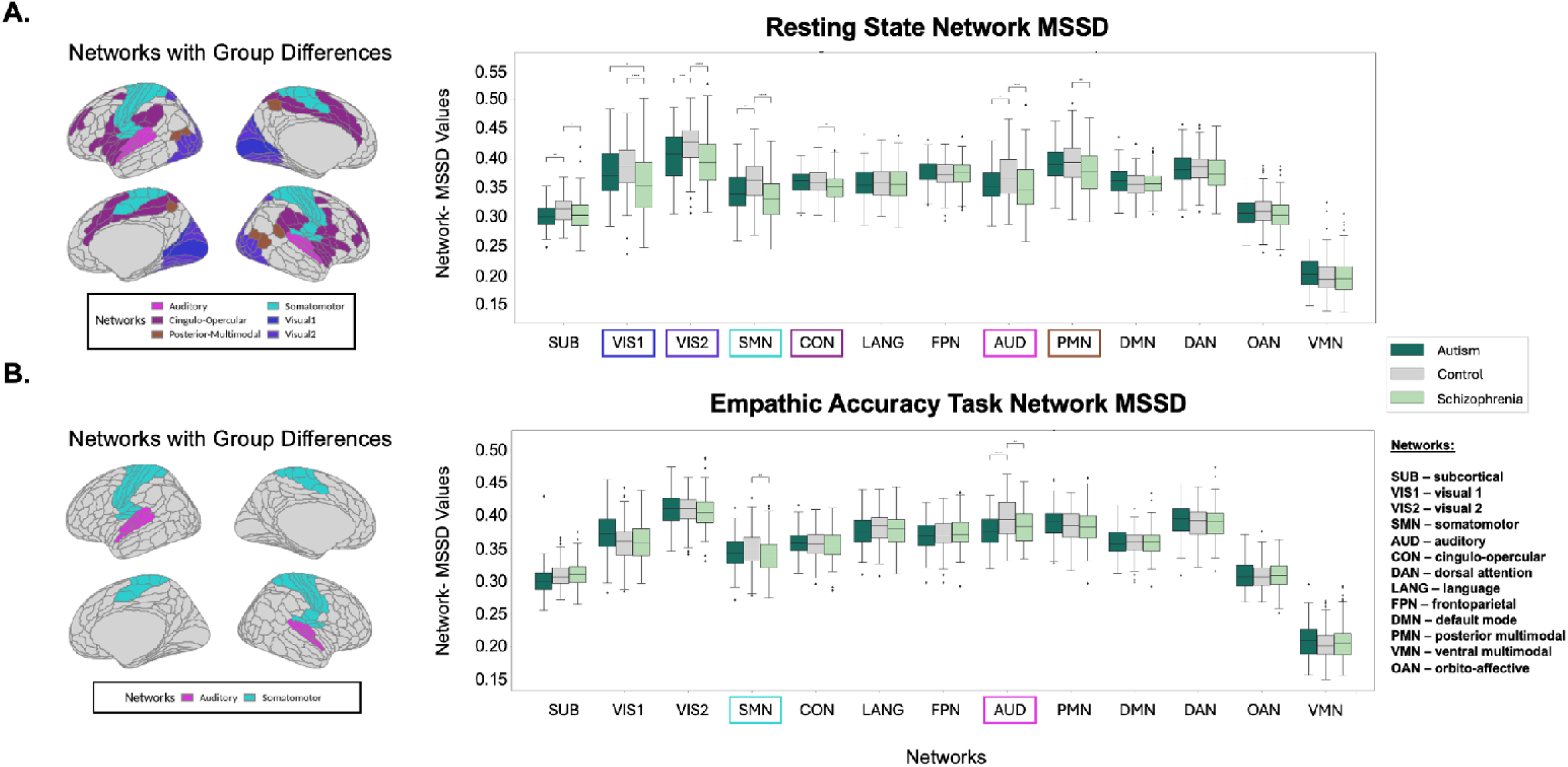
Network Variability via MSSD: **A)** Group-based differences in network variability, measured by mean squared successive difference (MSSD), during resting-state. Group differences were found across subcortical, visual 1, visual 2, somatomotor, cingulo-opercular, auditory, and posterior multimodal networks. **B)** Group-based differences in network variability, measured by MSSD, during the EA task. Group differences were found in the somatomotor and auditory networks.

During task (Fig. 3B; outlined in Table S4), TDC displayed greater MSSD compared to schizophrenia in the somatomotor and auditory networks. TDC displayed greater MSSD compared to the autism group in the auditory network. No significant differences were detected between autism and schizophrenia groups.

### 3.3 Network-Variability Regression Analysis with Behavioral/Cognitive/Clinical Variables of Interest

The relationship between network-based variability and social cognition, neurocognition, social functioning, symptom severity, and medication was examined across schizophrenia, autism, and TDC participants during rest and task across all 12 networks. No significant relationships were found between network MSSD and symptom severity (BPRS total score) during rest or task.

During rest, 20 significant relationships were detected after FDR correction between resting-state network MSSD and variables of interest (Fig. 4A). All relationships (excluding medication) were positively correlated, with significant relationships found in the visual 1, visual 2, somatomotor, auditory, cingulo-opercular, dorsal attention and subcortical networks across groups (outlined in Table S5). Broadly, greater network variability in visual 1, visual 2, somatomotor, auditory, cingulo-opercular, and dorsal attention networks was associated with higher MCCB scores, social cognition, and social functioning. When examining the relationship between medication and network variability, significant relationships were negatively correlated, indicating that lower MSSD predicted higher CPZ equivalents (i.e., higher antipsychotic medication dose) in the visual 1, somatomotor and auditory networks (Fig. 5 and Table S5).

**Figure 4.**
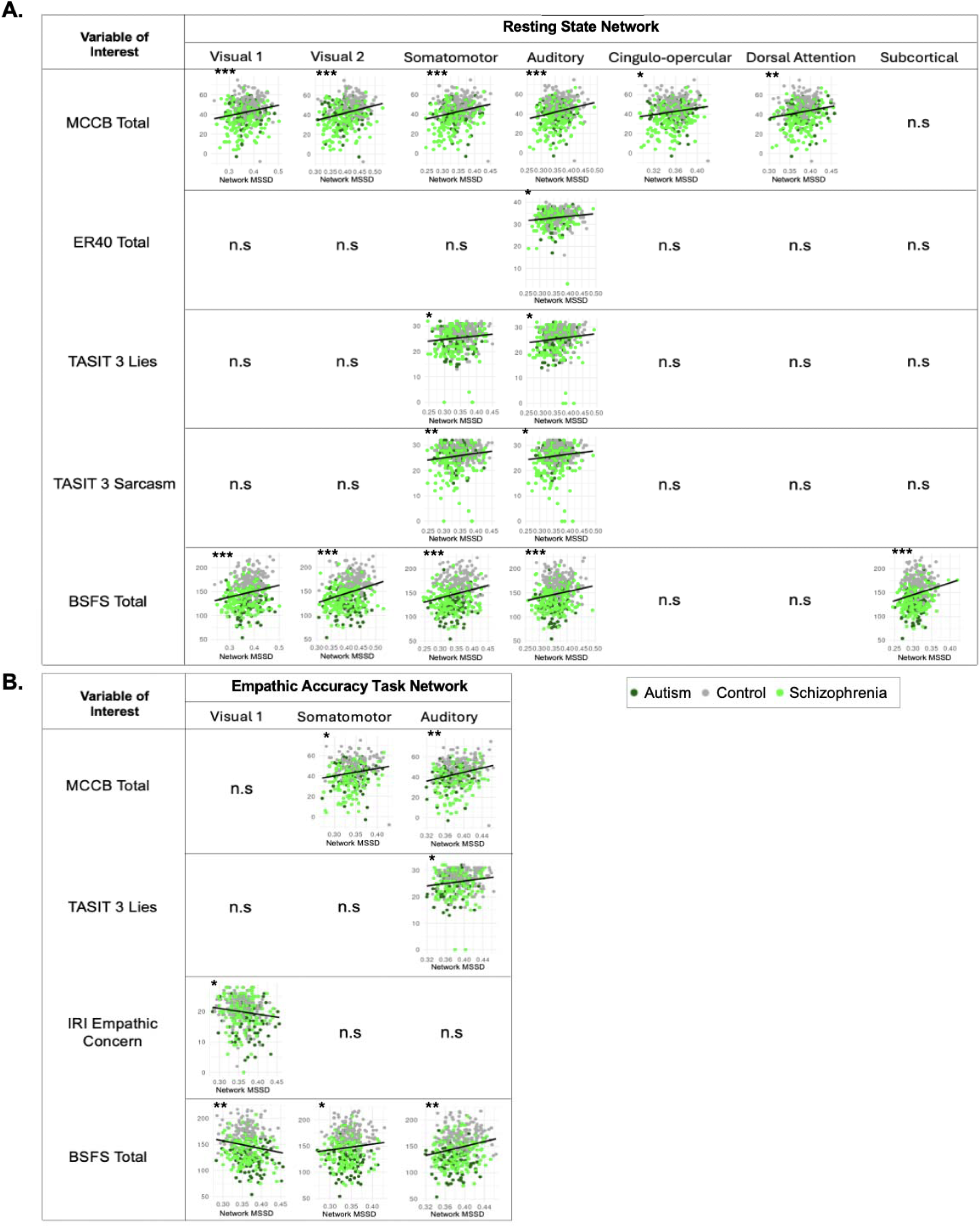
Relationship between Network Variability and Variables of Interest: Significant predictor variables and network variability during resting-state and EA task, where dots represent individual data points. Statistical significance is denoted by asterisks on each graph (*p < 0.05, **p < 0.01, ***p < 0.001). Only networks featuring at least one significant association are shown. **A)** Higher network variability during rest was positively correlated across variables of interest, including the BSFS total, MCCB composite, the TASIT3 Sarcasm & Lies, and ER40 total scores. **B)** Higher network variability during EA was positively correlated across variables of interest, including the BSFS total, MCCB composite, the TASIT3 Sarcasm & Lies, and IRI empathic concern scores. Negative correlations were found between visual 1 network variability and the BSFS total score and the IRI empathic concern score.

**Figure 5.**
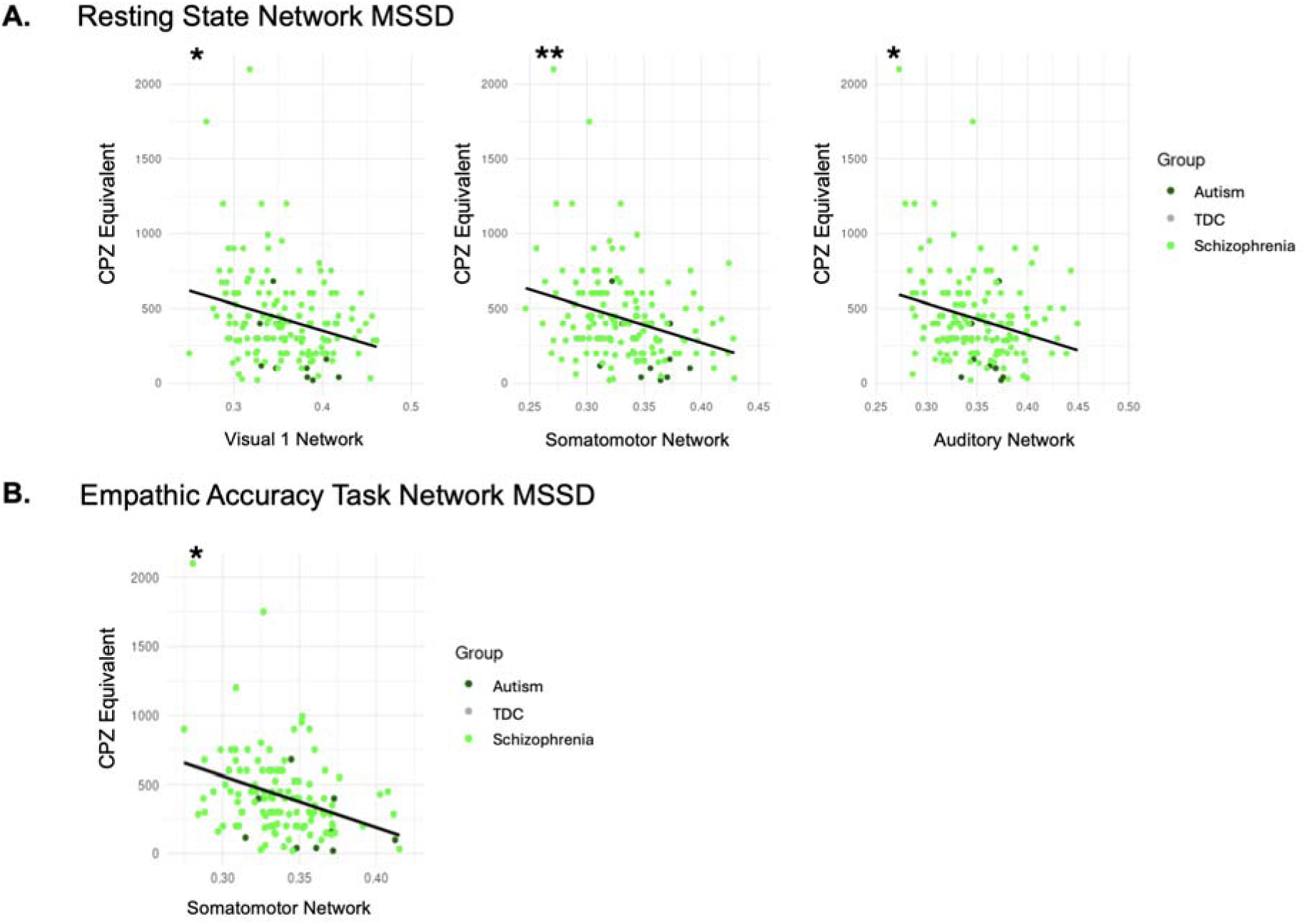
Relationship between Medication (CPZ equivalent) and Network Variability during Resting State and EA Task: Significant relationships between medication (CPZ equivalents) and network variability during resting state and the EA task, where dots represent individual data points. Statistical significance is denoted by asterisks on each graph (*p < 0.05, **p < 0.01) **A)** Network variability during rest was negatively correlated with medication across visual 1, somatomotor and auditory networks. **B)** Network variability during the EA task was negatively correlated with medication in the somatomotor network.

Plotted relationships and statistical outputs are outlined in Fig. 5 and Table S4 in the Supplemental Materials. Additionally, a group-by-network variability interaction term was also assessed to determine if relationships diverge across groups, with significant relationships found in the subcortical, somatomotor and cingulo-opercular networks (Fig. S3 and Table S6) and relationships with neurocognition and social functioning.

During the EA task, 8 significant relationships were detected between EA task network MSSD and our variables of interest (Fig. 4B; Table S7) across groups. Significant negative associations were found between MSSD within the visual 1 network and the BSFS total score (F(1, 369) = 10.57, p_FDR_ = 0.010) as well as the IRI empathic concern score (F(1, 369) = 7.31, p_FDR_ = 0.029), such that lower MSSD was associated with greater levels of self-reported social functioning as well as empathic concern. In contrast, significant positive associations were found between MSSD within the somatomotor and auditory networks, and our variables of interest (Fig. 4B). There was one significant relationship between medication and the visual 1 network during the EA task (F(1, 369) = 9.38, p_FDR_ = 0.011) (Fig. S3B and Table S7), in which the association was negatively correlated. A group-by-network variability interaction term was assessed but no significant relationships were found during task.

## 4. Discussion

In the present study, we investigated transdiagnostic intra-regional BOLD variability profiles derived during resting-state and EA task fMRI scanning in a large sample of individuals with diagnoses of schizophrenia or autism, and in TDC. Autism or schizophrenia displayed lower timeseries variability than TDC, both at rest and during task, predominantly in auditory, somatomotor, and visual regions/networks. Between-group differences (i.e., schizophrenia-TDC, autism-TDC or schizophrenia-autism) were more pronounced during resting-state than during the EA task, suggesting that resting-state variability may be more sensitive to diagnostic differences. Regression analyses examining relationships between network variability and cognitive/behavioral measures generally indicated that within visual, somatomotor, auditory, cingulo-opercular and dorsal attention networks, network-based variability was positively associated with social cognitive, neurocognitive, and social functioning scores across our transdiagnostic sample. These findings suggest that greater resting-state network variability is generally associated with better cognition and functional ability.

BOLD signal variability reflects dynamic neural processes [72] and is increasingly being recognized as an indicator of neural adaptability or flexibility [71]. Greater brain signal variability reflects a broader neural dynamic range, supporting more accurate stimulus processing [73], and provides a marker of overall cognitive efficiency [22]. In both resting-state and the EA task, participants with either autism or schizophrenia exhibited reduced regional and network-based variability compared to TDC, specifically in the visual, auditory, and somatomotor networks, and related sensory processing regions. These reductions were associated with lower social cognitive and neurocognitive performance, suggesting that decreased neural flexibility may contribute to cognitive difficulties in these populations. Both autism and schizophrenia groups feature atypicalities in sensory and information processing, and strong genetic heritability [74, 75]. Decreased BOLD signal variability across sensory processing regions may reflect decreased neural flexibility and abnormal sensory-motor processing that may cascade into cognitive deficits [76, 77], and may predict functional outcome [78]. Early abnormalities in sensory processing, particularly in auditory and multisensory integration, may compound over time, setting a maladaptive developmental trajectory that impairs the brain’s ability to detect and prioritize salient social cues. These disruptions can limit language acquisition, hinder social engagement, contribute to cascading impairments in cognitive domains across development, and ultimately to functional impacts [76].

Overall neurobiological patterns were largely overlapping between autism and schizophrenia, with less than 5% of regions exhibiting differences in timeseries variability between the groups (with differences only in the visual 1 network). This overlap is consistent with studies showing common biological differences in both conditions, including in genetic [79, 80] and neuroimaging studies [12, 81]. Our findings also align with recent work suggesting that social cognitive task-related activity and connectivity in schizophrenia and autism exist along a continuum with TDC [7, 35–38]. Reduced variability within sensory regions is consistent with findings such as reduced auditory steady-state response (ASSR) in both autism and schizophrenia. Both schizophrenia and autism have shown reduced gamma-band power and synchronization during ASSR, suggesting GABAergic and glutamatergic dysfunction [82, 83], related to sensory responses and cognitive functions like working memory [84, 85]. Decreased timeseries variability may also be exacerbated by medication effects in schizophrenia, as suggested by the negative relationships between visual 1, somatomotor and auditory network and CPZ equivalents (Fig. S3).

Across clinical groups, no significant group differences emerged in heteromodal networks such as the default mode, frontoparietal, and dorsal attention networks. Regions with higher BOLD signal variability often exhibit stronger functional connectivity and greater moment-to-moment synchrony with other regions, suggesting they are more functionally embedded within large-scale networks and play a central role in network integration [20, 86]. Recent findings from our group revealed decreased fractional amplitude of low-frequency fluctuations (fALFF) in visual regions across both autism and schizophrenia cohorts at rest, suggesting reduced spontaneous activity and signal variability in early sensory processing areas [87]. Consistent with this, prior studies have reported sensorimotor hypoconnectivity in both autism and schizophrenia, pointing to disruptions in sensory-motor integration [88–91].

Supporting this pattern, Fu et al. reported that both autism and schizophrenia groups tend to remain in connectivity states marked by weak intra-network coherence, particularly within sensorimotor systems, reflecting a generalized pattern of reduced and less dynamic functional integration [92]. This disruption was more pronounced in schizophrenia. Collectively, these findings suggest that individuals with autism and schizophrenia may possess more rigid, less functionally integrated sensorimotor networks, which could underlie shared impairments in social cognition and broader neurocognitive functioning.

Consistent with our previous work [35], group differences in variability were more pronounced at rest than during EA task performance. Task-evoked activity has been suggested to globally quench neural variability and reduce inter-regional cortical correlations [93], while resting-state activity is characterized by mind wandering and spontaneous, internally directed cognitive processes, which may in some cases amplify diagnostic differences in neural functioning [94, 95]. The EA task may engage neural strategies underlying emotional processing, which leads to more constrained social cognitive brain states [35, 37, 96]. While most associations indicated greater social cognitive resonance with higher variability, variability in the visual 1 network during EA showed negative associations with empathic concern and social functioning scores. This suggests excessive variability may be maladaptive in specific social cognitive contexts.

Several limitations of our study should be noted. First, while motion was controlled for at multiple stages of preprocessing and statistical analysis, it remains a potential confound in fMRI studies [97, 98], particularly in clinical populations. Second, medication effects, particularly in the schizophrenia group, may contribute to reduced variability, as suggested by the observed negative associations between CPZ equivalents and MSSD in key networks. However, it is important to note that the autism group also showed reduced variability in these networks despite low antipsychotic use, suggesting that this effect may persist even when controlling for CPZ equivalents. It is unclear if the associations between CPZ and MSSD are driven by medication effects, or if higher CPZ is a proxy for more severe illness. While the data acquisition between SPINS and SPIN-ASD were well harmonized and allowed for a large sample, participants with schizophrenia were recruited across multiple sites while autistic participants were drawn from a single site, resulting in differences in sample size and demographic variation between groups.

While samples were age-range matched and age, sex and mean FD were included as covariates in our analyses, the possibility of residual confounding from these or other unmeasured factors remains possible. Both the autism and schizophrenia samples included participants with relatively intact cognitive abilities (IQ scores for autism and schizophrenia samples were within the average range) who were able to complete neuropsychological assessments and brain imaging. Given the high prevalence of co-occurring intellectual disability, particularly in autism, this may limit the generalizability of our findings to the broader clinical population [99].

## 5. Conclusion

To our knowledge, this study was the first to conduct a transdiagnostic investigation of BOLD signal variability during resting-state and social cognitive task-based fMRI in schizophrenia and autism participants. Increased network-based variability was broadly linked to better social cognitive, neurocognitive and self-reported social functioning, reinforcing prior work hypothesizing that greater neural variability reflects a more flexible and efficient brain system [78, 100]. These results suggest that BOLD signal variability may serve as a sensitive and dynamic neural marker of individual differences in cognitive capacity, complementing traditional mean-based metrics [21]. Given its strong associations with social cognition, cognitive flexibility, and large-scale network function, brain signal variability emerges as a promising yet underutilized target for understanding neurobiological mechanisms in clinical populations. As a marker of neural flexibility, it provides a unique advantage in detecting local functional dysregulation and rigidity across disorders. Leveraging this measure in autism and schizophrenia may help uncover transdiagnostic mechanisms underlying shared cognitive characteristics, positioning BOLD variability as a potential biomarker for neural flexibility and a valuable target for future clinical interventions.

## Acknowledgements

We express our sincere gratitude to all individuals who contributed to this project in various capacities. We especially thank the study participants for their cooperation and patience. We also acknowledge the valuable support provided by the research staff involved in data collection, processing, and management. Their dedication and effort were instrumental to the successful completion of this work.

## References

1. Lord C, Elsabbagh M, Baird G, Veenstra-Vanderweele J. Autism spectrum disorder. Lancet. 2018;392:508–520.

2. Crespi B, Badcock C. Psychosis and autism as diametrical disorders of the social brain. Behav Brain Sci. 2008;31:241–261; discussion 261–320.

3. Sasson NJ, Morrison KE, Kelsven S, Pinkham AE. Social cognition as a predictor of functional and social skills in autistic adults without intellectual disability. Autism Res. 2020;13:259–270.

4. Halverson TF, Orleans-Pobee M, Merritt C, Sheeran P, Fett A-K, Penn DL. Pathways to functional outcomes in schizophrenia spectrum disorders: Meta-analysis of social cognitive and neurocognitive predictors. Neurosci Biobehav Rev. 2019;105:212–219.

5. Oliver LD, Moxon-Emre I, Lai M-C, Grennan L, Voineskos AN, Ameis SH. Social Cognitive Performance in Schizophrenia Spectrum Disorders Compared With Autism Spectrum Disorder: A Systematic Review, Meta-analysis, and Meta-regression. JAMA Psychiatry. 2021;78:281–292.

6. Rashidi AG, Oliver LD, Moxon-Emre I, Hawco C, Dickie EW, Pan R, et al. Comparative Analysis of Social Cognitive and Neurocognitive Performance Across Autism and Schizophrenia Spectrum Disorders. Schizophr Bull. 2025:sbaf005.

7. Oliver LD, Moxon-Emre I, Hawco C, Dickie EW, Dakli A, Lyon RE, et al. Task-based functional neural correlates of social cognition across autism and schizophrenia spectrum disorders. Molecular Autism. 2024;15.

8. Demetriou EA, DeMayo MM, Guastella AJ. Executive Function in Autism Spectrum Disorder: History, Theoretical Models, Empirical Findings, and Potential as an Endophenotype. Front Psychiatry. 2019;10:753.

9. Orellana G, Slachevsky A. Executive functioning in schizophrenia. Front Psychiatry. 2013;4:35.

10. McTeague LM, Huemer J, Carreon DM, Jiang Y, Eickhoff SB, Etkin A. Identification of Common Neural Circuit Disruptions in Cognitive Control Across Psychiatric Disorders. Am J Psychiatry. 2017;174:676–685.

11. Oliver, Haltigan, Gold, Foussias. Lower-and higher-level social cognitive factors across individuals with schizophrenia spectrum disorders and healthy controls: relationship with neurocognition and…. Schizophr Res. 2019. 2019.

12. Pinkham AE, Hopfinger JB, Pelphrey KA, Piven J, Penn DL. Neural bases for impaired social cognition in schizophrenia and autism spectrum disorders. Schizophr Res. 2008;99:164–175.

13. Ciaramidaro A, Bölte S, Schlitt S, Hainz D, Poustka F, Weber B, et al. Schizophrenia and autism as contrasting minds: neural evidence for the hypo-hyper-intentionality hypothesis. Schizophr Bull. 2015;41:171–179.

14. Hyatt CJ, Wexler BE, Pittman B, Nicholson A, Pearlson GD, Corbera S, et al. Atypical dynamic functional network connectivity state engagement during social-emotional processing in schizophrenia and autism. Cereb Cortex. 2022;32:3406–3422.

15. Hyatt CJ, Calhoun VD, Pittman B, Corbera S, Bell MD, Rabany L, et al. Default mode network modulation by mentalizing in young adults with autism spectrum disorder or schizophrenia. NeuroImage Clin. 2020;27:102343.

16. Hajdúk M, Pinkham AE, Penn DL, Harvey PD, Sasson NJ. Heterogeneity of social cognitive performance in autism and schizophrenia. Autism Res. 2022;15:1522–1534.

17. Mastrovito D, Hanson C, Hanson SJ. Differences in atypical resting-state effective connectivity distinguish autism from schizophrenia. Neuroimage Clin. 2018;18:367–376.

18. Logothetis NK, Wandell BA. Interpreting the BOLD signal. Annu Rev Physiol. 2004;66:735–769.

19. Uddin LQ. Bring the Noise: Reconceptualizing Spontaneous Neural Activity. Trends Cogn Sci. 2020;24:734–746.

20. Baracchini G, Mišić B, Setton R, Mwilambwe-Tshilobo L, Girn M, Nomi JS, et al. Inter-regional BOLD signal variability is an organizational feature of functional brain networks. Neuroimage. 2021;237:118149.

21. Garrett DD, Kovacevic N, McIntosh AR, Grady CL. Blood oxygen level-dependent signal variability is more than just noise. J Neurosci. 2010;30:4914–4921.

22. Garrett DD, Samanez-Larkin GR, MacDonald SWS, Lindenberger U, McIntosh AR, Grady CL. Moment-to-moment brain signal variability: a next frontier in human brain mapping? Neurosci Biobehav Rev. 2013;37:610–624.

23. Bryant AG, Aquino K, Parkes L, Fornito A, Fulcher BD. Extracting interpretable signatures of whole-brain dynamics through systematic comparison. PLoS Comput Biol. 2024;20:e1012692.

24. von Neumann J, Kent RH, Bellinson HR, Hart BI. The Mean Square Successive Difference. Ann Math Stat. 1941;12:153–162.

25. Nomi JS, Vij SG, Dajani DR, Steimke R, Damaraju E, Rachakonda S, et al. Chronnectomic patterns and neural flexibility underlie executive function. Neuroimage. 2017;147:861–871.

26. Goodman ZT, Nomi JS, Kornfeld S, Bolt T, Saumure RA, Romero C, et al. Brain signal variability and executive functions across the life span. Netw Neurosci. 2024;8:226–240.

27. Nomi JS, Schettini E, Voorhies W, Bolt TS, Heller AS, Uddin LQ. Resting-state brain signal variability in prefrontal cortex is associated with ADHD symptom severity in children. Front Hum Neurosci. 2018;12:90.

28. Easson AK, McIntosh AR. BOLD signal variability and complexity in children and adolescents with and without autism spectrum disorder. Dev Cogn Neurosci. 2019;36:100630.

29. Zhang Y, Yang R, Cai X. Frequency-specific alternations in the moment-to-moment BOLD signals variability in schizophrenia. Brain Imaging Behav. 2021;15:68–75.

30. Servaas MN, Kos C, Gravel N, Renken RJ, Marsman J-BC, van Tol M-J, et al. Rigidity in motor behavior and brain functioning in patients with schizophrenia and high levels of apathy. Schizophr Bull. 2019;45:542–551.

31. Sigar P, Kathrein N, Gragas E, Kupis L, Uddin LQ, Nomi JS. Age-related changes in brain signal variability in autism spectrum disorder. Mol Autism. 2025;16:8.

32. Baracchini G, Zhou Y, da Silva Castanheira J, Hansen JY, Rieck J, Turner GR, et al. The biological role of local and global fMRI BOLD signal variability in human brain organization. bioRxivorg. 2023.

33. Steinberg SN, King TZ. Within-individual BOLD signal variability and its implications for task-based cognition: A systematic review. Neuropsychol Rev. 2024;34:1115–1164.

34. Tsikonofilos K, Kumar A, Ampatzis K, Garrett DD, Månsson KNT. The promise of investigating neural variability in psychiatric disorders. Biol Psychiatry. 2025. 13 February 2025. 10.1016/j.biopsych.2025.02.004.

35. Secara MT, Oliver LD, Gallucci J, Dickie EW, Foussias G, Gold J, et al. Heterogeneity in functional connectivity: Dimensional predictors of individual variability during rest and task fMRI in psychosis. Prog Neuropsychopharmacol Biol Psychiatry. 2024;132:110991.

36. Viviano JD, Buchanan RW, Calarco N, Gold JM, Foussias G, Bhagwat N, et al. Resting-State Connectivity Biomarkers of Cognitive Performance and Social Function in Individuals With Schizophrenia Spectrum Disorder and Healthy Control Subjects. Biological Psychiatry. 2018;84:665–674.

37. Oliver LD, Hawco C, Homan P, Lee J, Green MF, Gold JM, et al. Social Cognitive Networks and Social Cognitive Performance Across Individuals With Schizophrenia Spectrum Disorders and Healthy Control Participants. Biol Psychiatry Cogn Neurosci Neuroimaging. 2021;6:1202–1214.

38. Hawco C, Buchanan RW, Calarco N, Mulsant BH, Viviano JD, Dickie EW, et al. Separable and Replicable Neural Strategies During Social Brain Function in People With and Without Severe Mental Illness. Am J Psychiatry. 2019;176:521–530.

39. Overall JE, Gorham DR. The Brief Psychiatric Rating Scale. Psychol Rep. 1962;10:799–812.

40. Buchanan RW, Javitt DC, Marder SR, Schooler NR, Gold JM, McMahon RP, et al. The Cognitive and Negative Symptoms in Schizophrenia Trial (CONSIST): the efficacy of glutamatergic agents for negative symptoms and cognitive impairments. Am J Psychiatry. 2007;164:1593–1602.

41. Birchwood M, Smith J, Cochrane R, Wetton S, Copestake S. The Social Functioning Scale. The development and validation of a new scale of social adjustment for use in family intervention programmes with schizophrenic patients. Br J Psychiatry. 1990;157:853–859.

42. Wechsler D. Wechsler Test of Adult Reading: WTAR. Psychological Corporation; 2001.

43. Wechsler D. Wechsler abbreviated scale of intelligence--second edition. PsycTESTS Dataset. 2018.

44. Leucht S, Samara M, Heres S, Davis JM. Dose Equivalents for Antipsychotic Drugs: The DDD Method. Schizophr Bull. 2016;42 Suppl 1:S90–S94.

45. Pinkham AE, Penn DL, Green MF, Buck B, Healey K, Harvey PD. The social cognition psychometric evaluation study: results of the expert survey and RAND panel. Schizophr Bull. 2014;40:813–823.

46. Morrison KE, Pinkham AE, Kelsven S, Ludwig K, Penn DL, Sasson NJ. Psychometric evaluation of social cognitive measures for adults with autism. Autism Res. 2019;12:766–778.

47. Kohler CG, Bilker W, Hagendoorn M, Gur RE, Gur RC. Emotion recognition deficit in schizophrenia: association with symptomatology and cognition. Biol Psychiatry. 2000;48:127–136.

48. Baron-Cohen S, Wheelwright S, Hill J, Raste Y, Plumb I. The ‘Reading the Mind in the Eyes’ Test Revised Version: A Study with Normal Adults, and Adults with Asperger Syndrome or High-functioning Autism. J Child Psychol Psychiatry. 2001;42:241–251.

49. McDonald S, Flanagan S, Rollins J. The awareness of social inference test (revised). 2011.

50. Davis MH. Measuring individual differences in empathy: Evidence for a multidimensional approach. J Pers Soc Psychol. 1983;44:113–126.

51. Shamay-Tsoory SG, Aharon-Peretz J, Perry D. Two systems for empathy: a double dissociation between emotional and cognitive empathy in inferior frontal gyrus versus ventromedial prefrontal lesions. Brain. 2009;132:617–627.

52. Zaki J, Weber J, Bolger N, Ochsner K. The neural bases of empathic accuracy. Proc Natl Acad Sci U S A. 2009;106:11382–11387.

53. Kern RS, Penn DL, Lee J, Horan WP, Reise SP, Ochsner KN, et al. Adapting social neuroscience measures for schizophrenia clinical trials, Part 2: trolling the depths of psychometric properties. Schizophr Bull. 2013;39:1201–1210.

54. Olbert CM, Penn DL, Kern RS, Lee J, Horan WP, Reise SP, et al. Adapting social neuroscience measures for schizophrenia clinical trials, part 3: fathoming external validity. Schizophr Bull. 2013;39:1211–1218.

55. Nuechterlein KH, Green MF, Kern RS, Baade LE, Barch DM, Cohen JD, et al. The MATRICS Consensus Cognitive Battery, part 1: test selection, reliability, and validity. Am J Psychiatry. 2008;165:203–213.

56. Ludwig KA, Pinkham AE, Harvey PD, Kelsven S, Penn DL. Social cognition psychometric evaluation (SCOPE) in people with early psychosis: A preliminary study. Schizophr Res. 2017;190:136–143.

57. Kuo SS, Wojtalik JA, Mesholam-Gately RI, Keshavan MS, Eack SM. Transdiagnostic validity of the MATRICS Consensus Cognitive Battery across the autism-schizophrenia spectrum. Psychol Med. 2020;50:1623–1632.

58. Esteban O, Markiewicz CJ, Blair RW, Moodie CA, Isik AI, Erramuzpe A, et al. fMRIPrep: a robust preprocessing pipeline for functional MRI. Nat Methods. 2019;16:111–116.

59. Gorgolewski KJ, Esteban O, Markiewicz CJ, Ziegler E. Nipype. Softw Pract Exp. 2018. 2018.

60. Dickie EW, Anticevic A, Smith DE, Coalson TS, Manogaran M, Calarco N, et al. Ciftify: A framework for surface-based analysis of legacy MR acquisitions. Neuroimage. 2019;197:818–826.

61. Muschelli J, Nebel MB, Caffo BS, Barber AD, Pekar JJ, Mostofsky SH. Reduction of motion-related artifacts in resting state fMRI using aCompCor. Neuroimage. 2014;96:22–35.

62. Satterthwaite TD, Elliott MA, Gerraty RT, Ruparel K, Loughead J, Calkins ME, et al. An improved framework for confound regression and filtering for control of motion artifact in the preprocessing of resting-state functional connectivity data. Neuroimage. 2013;64:240–256.

63. Glasser MF, Coalson TS, Robinson EC, Hacker CD, Harwell J, Yacoub E, et al. A multi-modal parcellation of human cerebral cortex. Nature. 2016;536:171–178.

64. Ji JL, Spronk M, Kulkarni K, Repovš G, Anticevic A, Cole MW. Mapping the human brain’s cortical-subcortical functional network organization. Neuroimage. 2019;185:35–57.

65. Tian Y, Margulies DS, Breakspear M, Zalesky A. Topographic organization of the human subcortex unveiled with functional connectivity gradients. Nat Neurosci. 2020;23:1421–1432.

66. Garrett DD, Kovacevic N, McIntosh AR, Grady CL. The Importance of Being Variable. J Neurosci. 2011;31:4496–4503.

67. Wehrheim MH, Faskowitz J, Schubert A-L, Fiebach CJ. Reliability of variability and complexity measures for task and task-free BOLD fMRI. Hum Brain Mapp. 2024;45:e26778.

68. Johnson WE, Li C, Rabinovic A. Adjusting batch effects in microarray expression data using empirical Bayes methods. Biostatistics. 2007;8:118–127.

69. Fortin J-P, Cullen N, Sheline YI, Taylor WD, Aselcioglu I, Cook PA, et al. Harmonization of cortical thickness measurements across scanners and sites. Neuroimage. 2018;167:104–120.

70. Mancuso F, Horan WP, Kern RS, Green MF. Social cognition in psychosis: multidimensional structure, clinical correlates, and relationship with functional outcome. Schizophr Res. 2011;125:143–151.

71. Zhang C, Beste C, Prochazkova L, Wang K, Speer SPH, Smidts A, et al. Resting-state BOLD signal variability is associated with individual differences in metacontrol. Sci Rep. 2022;12:18425.

72. Grady CL, Garrett DD. Understanding variability in the BOLD signal and why it matters for aging. Brain Imaging Behav. 2014;8:274–283.

73. Garrett DD, Epp SM, Kleemeyer M, Lindenberger U, Polk TA. Higher performers upregulate brain signal variability in response to more feature-rich visual input. Neuroimage. 2020;217:116836.

74. Sandin S, Lichtenstein P, Kuja-Halkola R, Hultman C, Larsson H, Reichenberg A. The heritability of autism spectrum disorder. JAMA. 2017;318:1182–1184.

75. Aukes MF, Alizadeh BZ, Sitskoorn MM, Selten J-P, Sinke RJ, Kemner C, et al. Finding suitable phenotypes for genetic studies of schizophrenia: heritability and segregation analysis. Biol Psychiatry. 2008;64:128–136.

76. Thye MD, Bednarz HM, Herringshaw AJ, Sartin EB, Kana RK. The impact of atypical sensory processing on social impairments in autism spectrum disorder. Dev Cogn Neurosci. 2018;29:151– 167.

77. Javitt DC, Freedman R. Sensory processing dysfunction in the personal experience and neuronal machinery of schizophrenia. Am J Psychiatry. 2015;172:17–31.

78. Månsson KNT, Waschke L, Manzouri A, Furmark T, Fischer H, Garrett DD. Moment-to-moment brain signal variability reliably predicts psychiatric treatment outcome. Biol Psychiatry. 2022;91:658–666.

79. Cross-Disorder Group of the Psychiatric Genomics Consortium. Identification of risk loci with shared effects on five major psychiatric disorders: a genome-wide analysis. Lancet. 2013;381:1371– 1379.

80. Geschwind DH, Flint J. Genetics and genomics of psychiatric disease. Science. 2015;349:1489– 1494.

81. Sugranyes G, Kyriakopoulos M, Corrigall R, Taylor E, Frangou S. Autism spectrum disorders and schizophrenia: meta-analysis of the neural correlates of social cognition. PLoS One. 2011;6:e25322.

82. Sugiyama S, Ohi K, Kuramitsu A, Takai K, Muto Y, Taniguchi T, et al. The auditory steady-state response: Electrophysiological index for sensory processing dysfunction in psychiatric disorders. Front Psychiatry. 2021;12:644541.

83. Gao R, Penzes P. Common mechanisms of excitatory and inhibitory imbalance in schizophrenia and autism spectrum disorders. Curr Mol Med. 2015;15:146–167.

84. Lundqvist M, Rose J, Herman P, Brincat SL, Buschman TJ, Miller EK. Gamma and beta bursts underlie working memory. Neuron. 2016;90:152–164.

85. Yamamoto J, Suh J, Takeuchi D, Tonegawa S. Successful execution of working memory linked to synchronized high-frequency gamma oscillations. Cell. 2014;157:845–857.

86. Garrett DD, Epp SM, Perry A, Lindenberger U. Local temporal variability reflects functional integration in the human brain. Neuroimage. 2018;183:776–787.

87. Bagheri S, Yu J-C, Gallucci J, Tan V, Oliver LD, Dickie EW, et al. Transdiagnostic neurobiology of social cognition and individual variability as measured by fractional amplitude of low-frequency fluctuation in autism and schizophrenia spectrum disorders. Biol Psychiatry Cogn Neurosci Neuroimaging. 2025. 21 April 2025. 10.1016/j.bpsc.2025.04.004.

88. Brandl F, Avram M, Weise B, Shang J, Simões B, Bertram T, et al. Specific Substantial Dysconnectivity in Schizophrenia: A Transdiagnostic Multimodal Meta-analysis of Resting-State Functional and Structural Magnetic Resonance Imaging Studies. Biol Psychiatry. 2019;85:573–583.

89. Dong D, Wang Y, Chang X, Luo C, Yao D. Dysfunction of Large-Scale Brain Networks in Schizophrenia: A Meta-analysis of Resting-State Functional Connectivity. Schizophr Bull. 2018;44:168–181.

90. Oldehinkel M, Mennes M, Marquand A, Charman T, Tillmann J, Ecker C, et al. Altered connectivity between cerebellum, visual, and sensory-motor networks in autism spectrum disorder: Results from the EU-AIMS Longitudinal European Autism Project. Biol Psychiatry Cogn Neurosci Neuroimaging. 2019;4:260–270.

91. Ilioska I, Oldehinkel M, Llera A, Chopra S, Looden T, Chauvin R, et al. Connectome-wide mega-analysis reveals robust patterns of atypical functional connectivity in autism. Biol Psychiatry. 2023;94:29–39.

92. Fu Z, Sui J, Turner JA, Du Y, Assaf M, Pearlson GD, et al. Dynamic functional network reconfiguration underlying the pathophysiology of schizophrenia and autism spectrum disorder. Hum Brain Mapp. 2021;42:80–94.

93. Ito T, Brincat SL, Siegel M, Mill RD, He BJ, Miller EK, et al. Task-evoked activity quenches neural correlations and variability across cortical areas. PLoS Comput Biol. 2020;16:e1007983.

94. Diaz BA, Van Der Sluis S, Moens S, Benjamins JS, Migliorati F, Stoffers D, et al. The Amsterdam Resting-State Questionnaire reveals multiple phenotypes of resting-state cognition. Front Hum Neurosci. 2013;7:446.

95. Marchetti A, Baglio F, Costantini I, Dipasquale O, Savazzi F, Nemni R, et al. Theory of Mind and the Whole Brain Functional Connectivity: Behavioral and Neural Evidences with the Amsterdam Resting State Questionnaire. Front Psychol. 2015;6:1855.

96. Cole MW, Bassett DS, Power JD, Braver TS, Petersen SE. Intrinsic and task-evoked network architectures of the human brain. Neuron. 2014;83:238–251.

97. Savalia NK, Agres PF, Chan MY, Feczko EJ, Kennedy KM, Wig GS. Motion-related artifacts in structural brain images revealed with independent estimates of in-scanner head motion. Hum Brain Mapp. 2017;38:472–492.

98. Van Dijk KRA, Sabuncu MR, Buckner RL. The influence of head motion on intrinsic functional connectivity MRI. Neuroimage. 2012;59:431–438.

99. Lai M-C, Lombardo MV, Baron-Cohen S. vol. 383, issue 9920. Autism Lancet. 2014:896–910.

100. Waschke L, Kloosterman NA, Obleser J, Garrett DD. Behavior needs neural variability. Neuron. 2021;109:751–766.

