## Supplemental_Information for "Transdiagnostic Profiles of BOLD Signal Variability in Autism and Schizophrenia Spectrum Disorders: Associations with Cognition and Functioning"

***Supplementary Information***

**Supplementary Methods and Materials**

**Participant Inclusion/Exclusion Criteria**

Clinical diagnoses were confirmed by a clinical team; participants with schizophrenia met DSM-5 criteria for schizophrenia, schizoaffective disorder, schizophreniform disorder, delusional disorder, or psychotic disorder not otherwise specified, assessed using the Structured Clinical Interview for DSM-IV-TR [[1]](https://paperpile.com/c/lepNMJ/XoEUq). Participants with autism met DSM-5 criteria for autism and were assessed using the Autism Diagnostic Observation Schedule 2nd Edition-Module 4 [[2]](https://paperpile.com/c/lepNMJ/HWcP). Information regarding participant medication was collected, and chlorpromazine (CPZ) equivalents were calculated for the schizophrenia and autism groups [[3]](https://paperpile.com/c/lepNMJ/XzC6R). Individuals with schizophrenia and autism were symptomatically stable, and had no change in antipsychotic medication or decrement in functioning/support level during the 30 days prior to enrollment. TDC did not have a current or past Axis I psychiatric disorder, excepting adjustment disorder, phobic disorder, and past major depressive disorder (over two years prior; presently unmedicated), or a first degree relative with a history of psychotic mental disorder. Additional exclusion criteria across all participants included: an IQ<70 (SPIN-ASD: estimated using the Wechsler Abbreviated Scale of Intelligence – Second Edition, WASI-II [[4]](https://paperpile.com/c/lepNMJ/Fck0); SPINS: estimated using the Wechsler Test of Adult Reading, WTAR [[5]](https://paperpile.com/c/lepNMJ/yZ2G)), a history of head trauma resulting in unconsciousness, metal implants or pacemakers, a substance use disorder (confirmed by urine toxicology screening), intellectual disability, type 1 diabetes, pregnancy, debilitating or unstable medical illness, recent epilepsy or stroke, or other neurological diseases, and non-English speakers.

**Empathic Accuracy (EA) Task**

The EA task was conducted during fMRI, with participants trained on a practice version of the task in a mock scanner beforehand. During the task, participants viewed nine videos depicting emotional autobiographical events (4 positive, 5 negative) with diverse adult actors, varying in age, race, and ethnicity (details in [[6, 7]](https://paperpile.com/c/lepNMJ/Xeme+oPgC)). EA scores were calculated by correlating participants' ratings with the self-ratings of the actors in the videos, and EA values were then Fisher r-to-z transformed. During the control condition (two 40-second interleaved videos per run), participants rated the lightness or darkness of a changing grayscale circle on a 9-point scale to ensure task engagement and comprehension. This condition is included to ensure that participants are engaged in the task (controlling for avolition) and comprehend it.

**MRI Data Acquisition**

MRI scans were collected using harmonized scanning parameters on 3T scanners with multichannel head coils, including a General Electric Discovery (N=115; CAMH) and Siemens Prisma (N=148; CAMH), a General Electric Signa (N=18; ZHH) and Siemens Prisma (N=55; ZHH), and a Siemens Tim Trio (N=30; MPRC) and Siemens Prisma (N=47; MPRC).

**MRI Exclusion Criteria**

All scans were quality checked by experienced research staff, using an in-house quality control system dashboard ([https://github.com/TIGRLab/dashboard)](https://github.com/TIGRLab/dashboard) prior to preprocessing, and after preprocessing [[8, 9]](https://paperpile.com/c/lepNMJ/xNxY+mt76). This included qualitative (e.g., detection of ghosting or ringing) and quantitative (e.g., framewise displacement) monitoring. Participants with excessive motion (mean FD per run > 0.5 mm) were removed during the EA task or RS scan, as well as EA or control task performance indicative of disengagement or a lack of comprehension. Specifically, participants were excluded if they had any EA videos with no responses, or more than one video with only one response, or a control task accuracy of < 20% and one or two responses in any EA video. Participants were also excluded from analyses if they were missing any cognitive metrics. See the consort flow diagram for inclusion/exclusion details (Figure S1).

**fMRI Preprocessing**

Anatomical T1-weighted images were corrected for intensity non-uniformity using ANTS 2.2.3

[[10]](https://paperpile.com/c/lepNMJ/iheW) and skull-stripping using Nipype implementation of the antsBrainExtraction.sh workflow (from ANTs). Brain tissue segmentation of cerebrospinal fluid, white-matter and gray-matter was performed using FSL 5.0.9 [[11]](https://paperpile.com/c/lepNMJ/Ugtv) Brain surfaces were reconstructed using FreeSurfer 6.0.1[[12]](https://paperpile.com/c/lepNMJ/gHpB). The extracted brain was spatially normalized to MNI space using nonlinear registration with ANTs[[10]](https://paperpile.com/c/lepNMJ/iheW). For each of the fMRI runs, fieldmap-less distortion correction was performed using fMRIPrep. Registration is performed using ANTs [[13]](https://paperpile.com/c/lepNMJ/ofp95). Functional data was coregistered to the corresponding T1-weighted image using Freesurfer’s boundary-based registration with six degrees of freedom. Head-motion parameters were estimated using MCFLIRT (FSL 5.0.9; [[14]](https://paperpile.com/c/lepNMJ/VAmhM)), and slice-time correction was performed using 3dTshift from AFNI [[15]](https://paperpile.com/c/lepNMJ/QWzZ) RRID_SCR_005927). The Ciftify toolbox ([[16]](https://paperpile.com/c/lepNMJ/flpM); [https://github.com/edickie/ciftify)](https://github.com/edickie/ciftify)) was used to transform the functional data onto the cortical surface (fsLR32k space; [[17]](https://paperpile.com/c/lepNMJ/bgcy9) using a non-linear transform to the MNI152 template via FSL's FNIRT. Four TRs were dropped for each EA scan and three TRs for the resting state scan, and data was smoothed at 2mm FWHM on the cortical surface. For resting state and EA, detrending, band-pass filtering (0.01-0.1 Hz), and nuisance regression was also performed using Ciftify. The nuisance regression model included regressors for the six head motion correction parameters, mean white matter signal, mean cerebral spinal fluid signal, the square, derivative, and square of the derivative for each of these regressors, and global brain signal (generated by fMRIPrep)[[18, 19]](https://paperpile.com/c/lepNMJ/g7Q0o+Qkumg).

**Statistical Analysis**

A Wilcoxon signed-rank test between each participant’s network-based MSSD values was run to determine if participants displayed differences in network timeseries variability during the EA task when compared to the same participants at rest.

**Supplementary Results**

**Network-Based Variability: MSSD Comparison Across EA Task and Rest**

Network-based MSSD was significantly higher during the EA task when compared to rest in the language (Wilcoxon signed-rank test; V = 59206, p_FDR_ = <2.2e-16), auditory (Wilcoxon signed-rank test; V = 57966, p_FDR_ = <2.2e-16), dorsal attention (Wilcoxon signed-rank test; V = 49420, p_FDR_ = 4.0e-16) and ventral multimodal networks (Wilcoxon signed-rank test; V = 42239, p_FDR_ = 8.0e-06); network MSSD was significantly higher during rest solely in the visual network (Wilcoxon signed-rank test; V = 26287, p_FDR_ = 9.9e-04) when compared to EA. Plotted distributions and statistical outputs are outlined in Figure S2 and Table S2.

**Figure S1: Consort Flow Diagram**

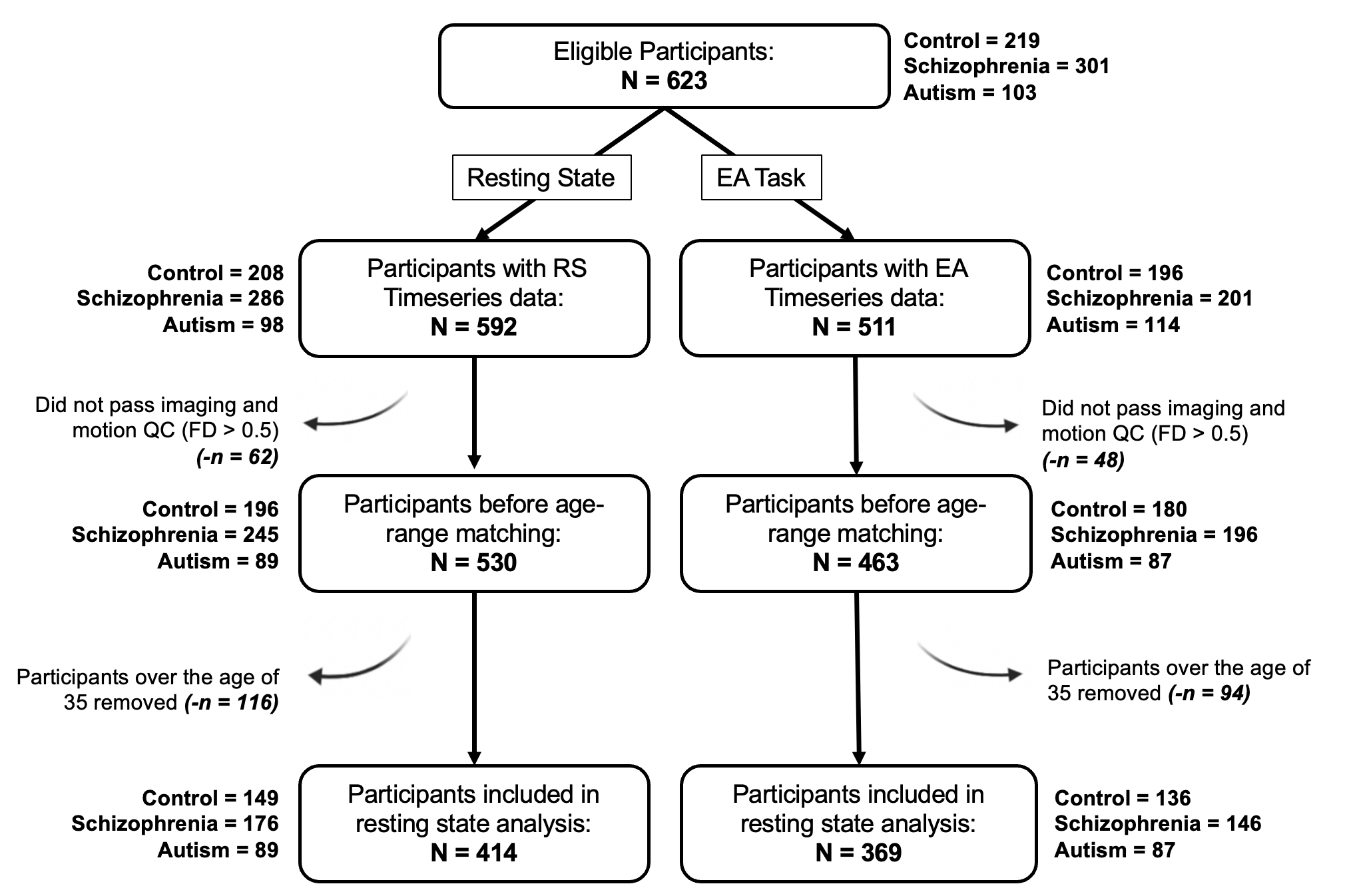
**Figure S1. Consort Flow Diagram:** Participant data was evaluated for eligibility based on a quality-control (QC) criteria. The initial data from 219 TDC, 301 schizophrenia and 114 autism (n = 634) participants were assessed for available MRI data; 42 participants were excluded from resting state and 123 the Empathic Accuracy (EA) task. Following this, data was controlled for excessive motion and resting state and EA task QC criteria; 62 participants were excluded during rest and 48 participants were excluded during the EA task. Lastly, participants over the age of 35 were excluded from the sample; 116 participants were removed from rest and 94 from the EA task. The resulting data from 149 TDC, 176 schizophrenia, 89 autism individuals with EA task, and 136 TDC, 146 schizophrenia, 87 autism individuals with RS data, were included for analysis.

**Table S1: Medication Usage Across Participants**

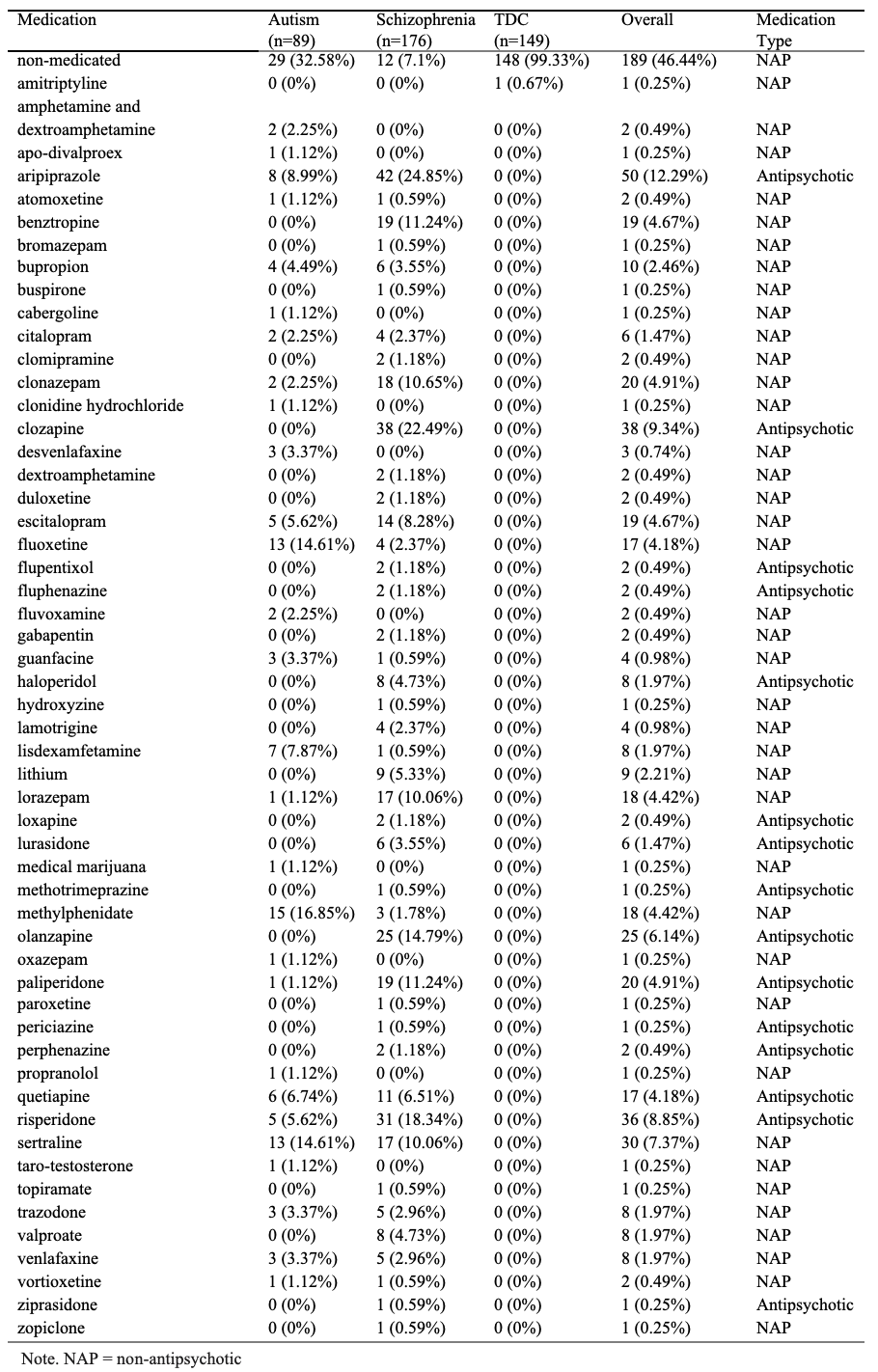

**Figure S2: Network-Based Variability: Plotted MSSD Value Distribution EA Task vs Rest**
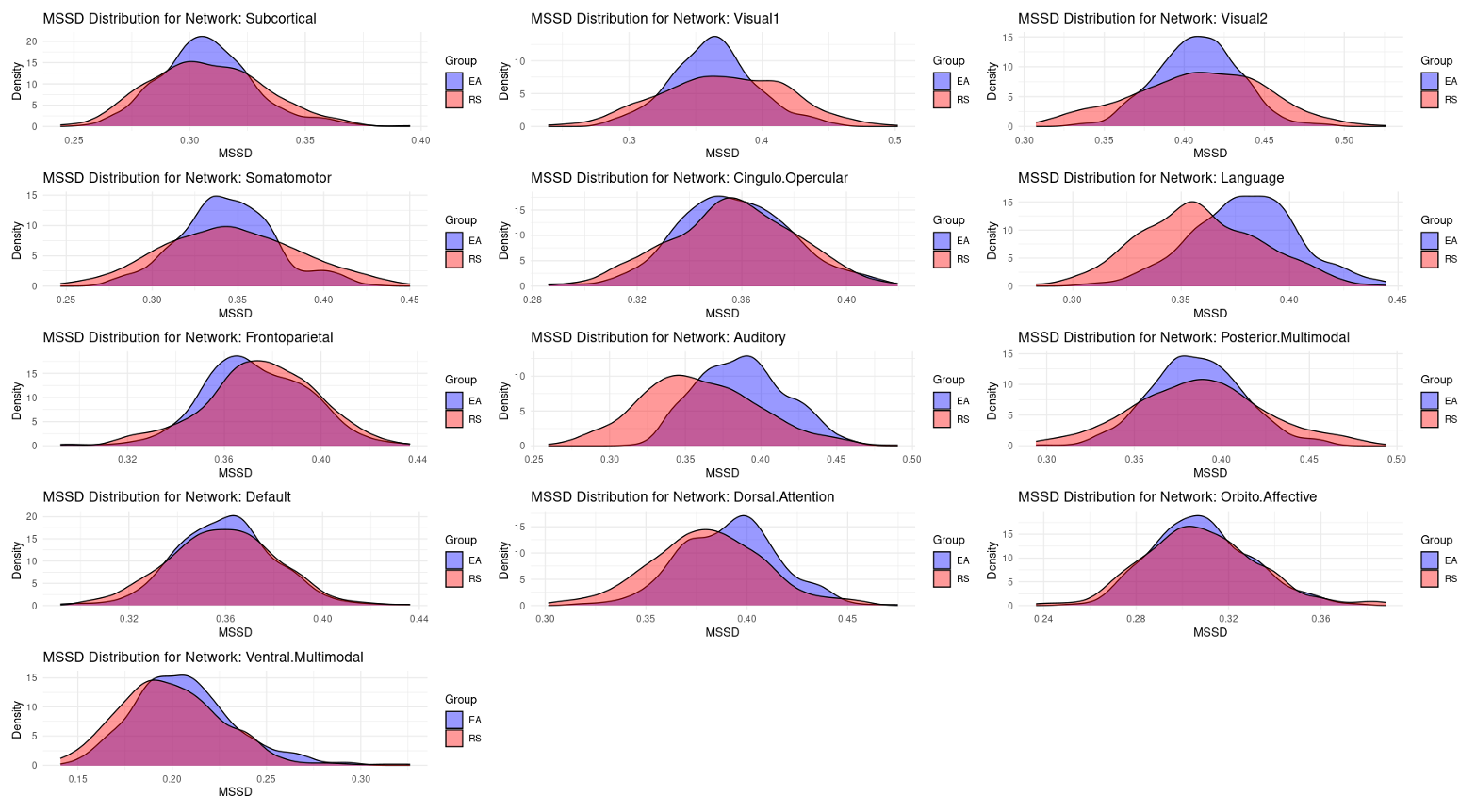

**Figure S2. Network-Based Variability: Plotted MSSD Value Distribution EA Task vs Rest:** Density plot depicting distribution of network-specific MSSD, comparing EA task (purple/blue) to resting state (orange) variability.

**Table S2: EA Task vs Resting State Network Variability**
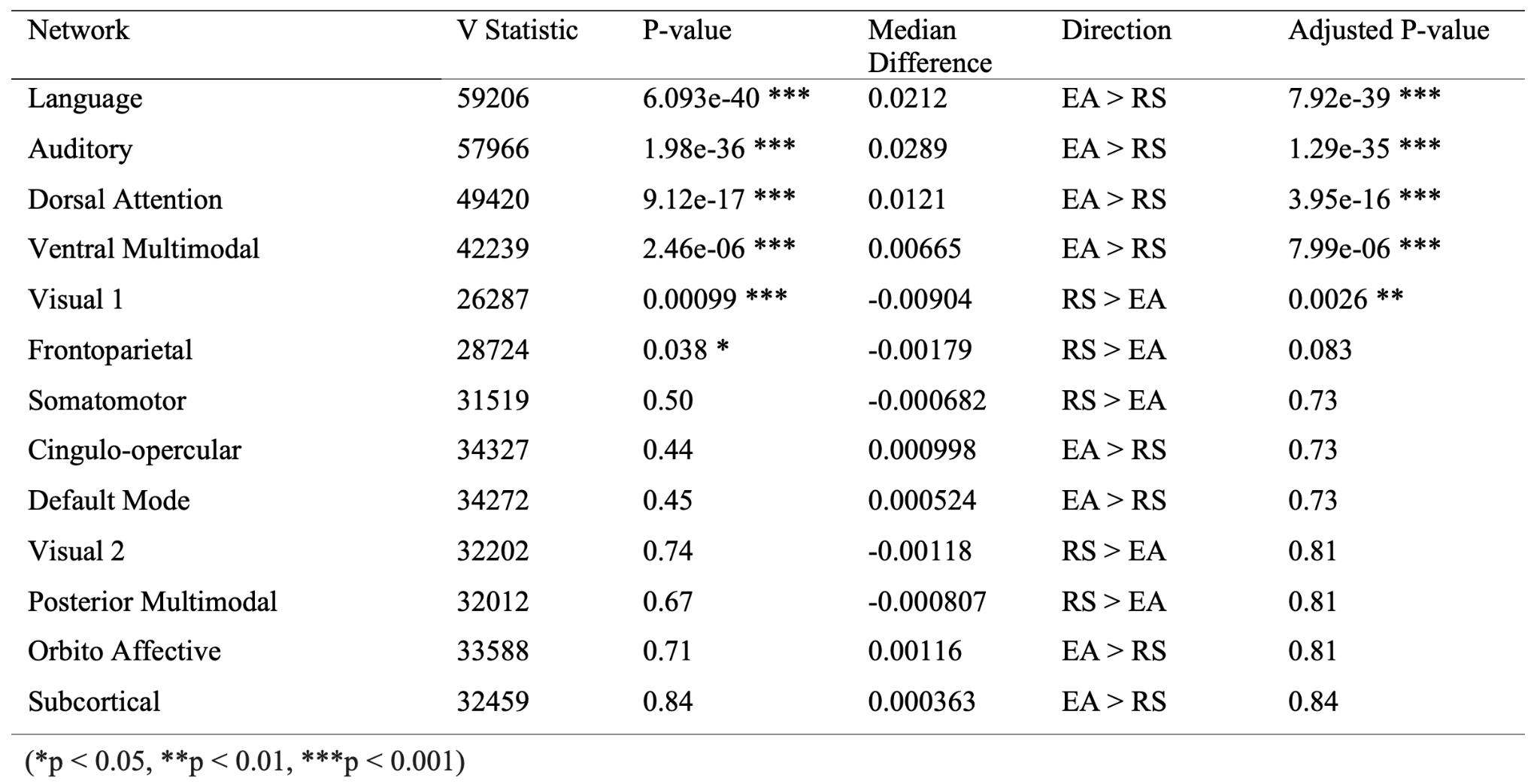

**Table S3: Group Comparison of Network Variability Differences during Resting State**
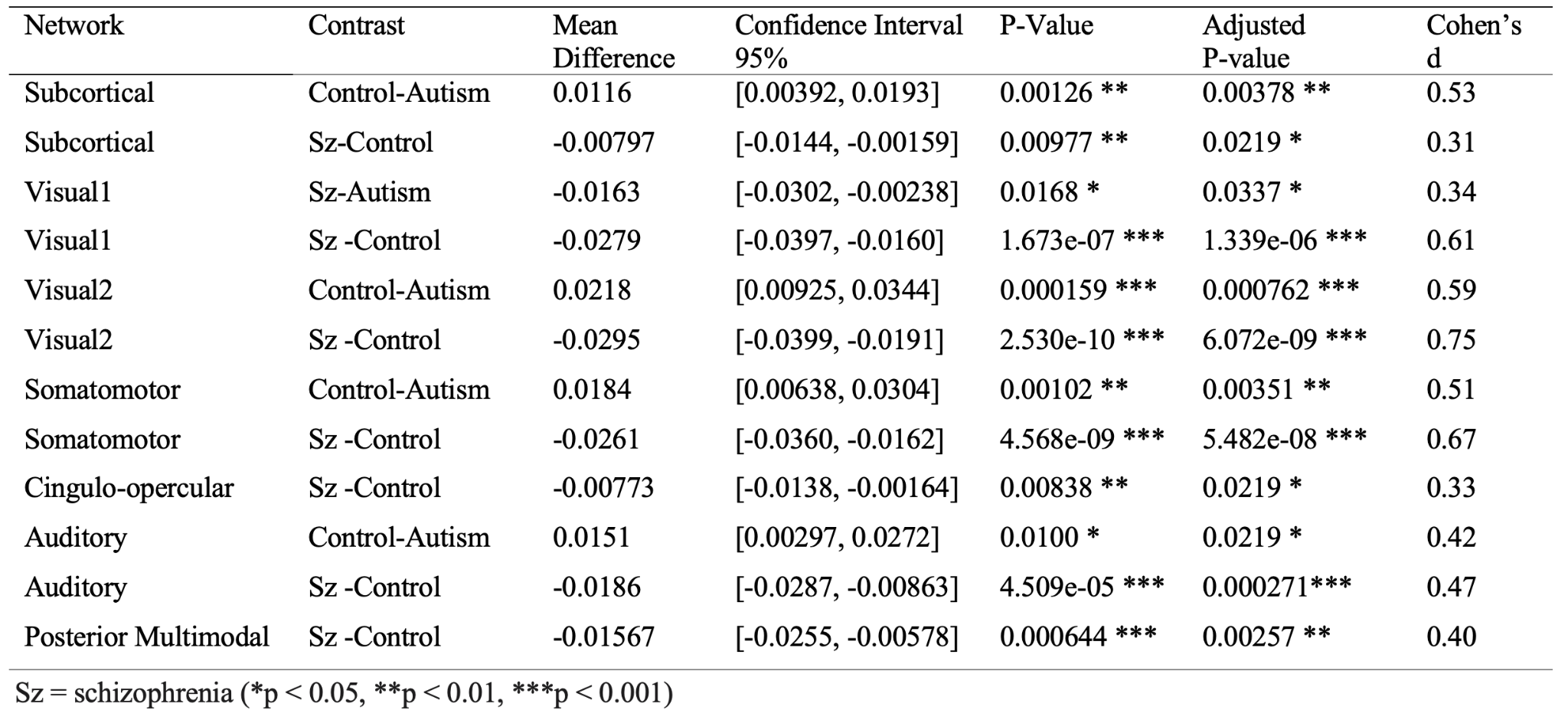

**Table S4: Group Comparison of Network Variability Differences during EA Task**

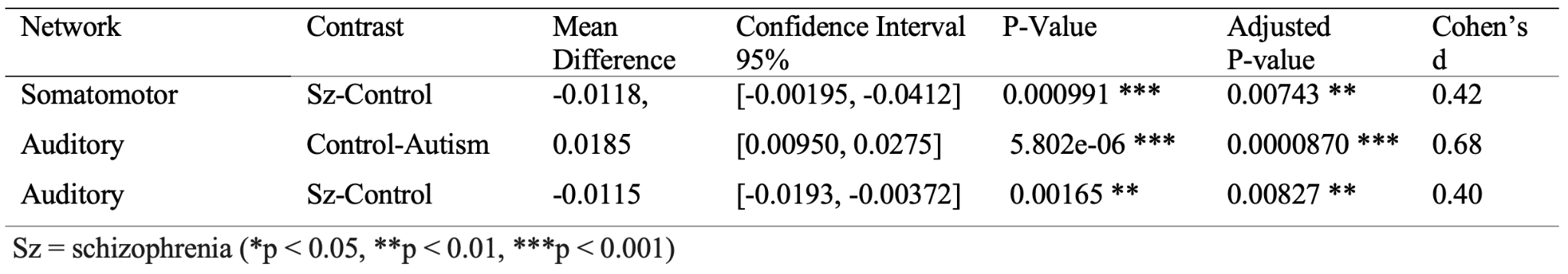
**Table S5: Statistical Outputs Assessing the relationship between Network-Variability and Variables of Interest during Resting State**

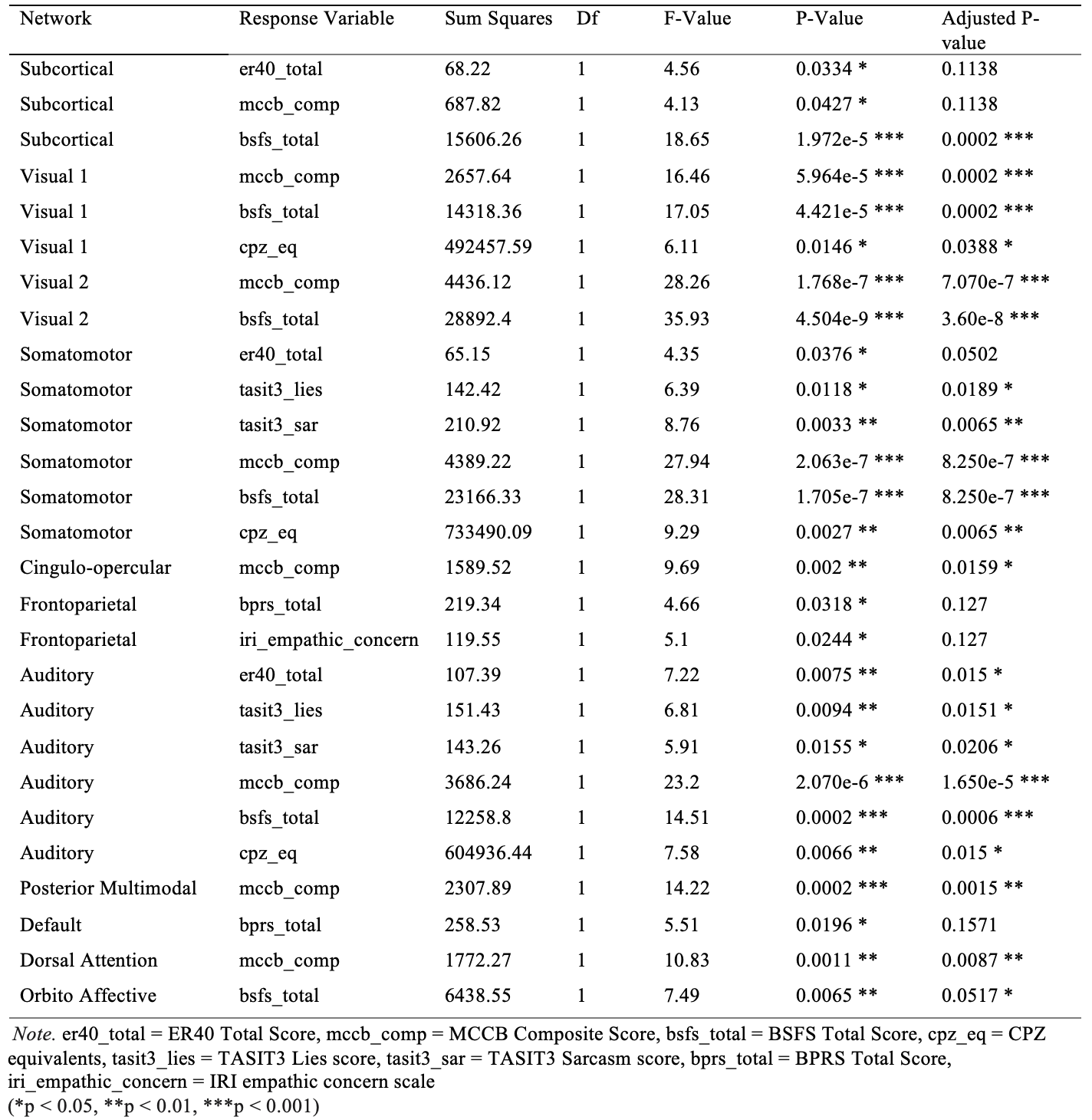

**Figure S3: Relationship between Variables of Interest and Network Variability during**

**Resting State + Group Interaction Terms**

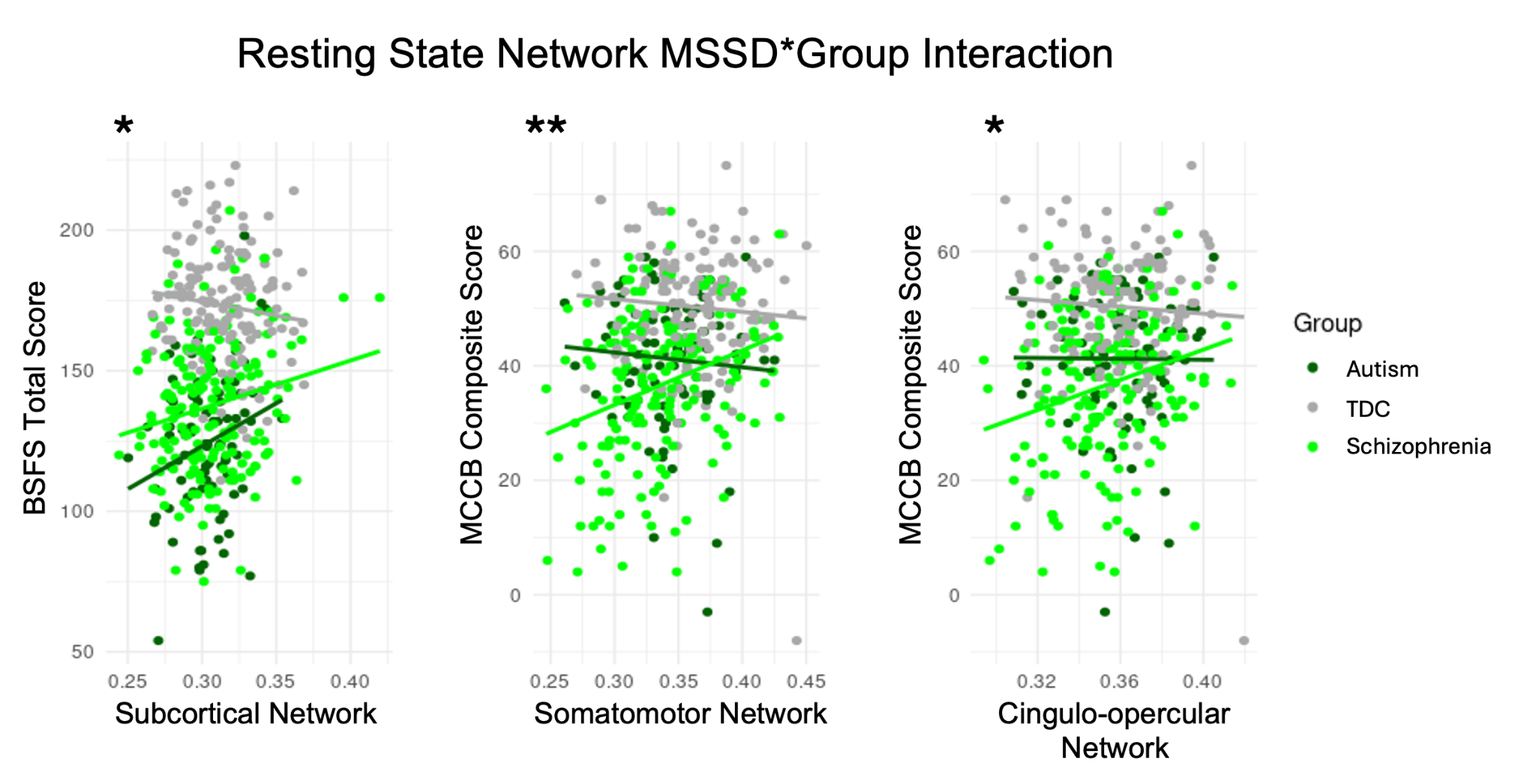

**Figure S3: Relationship between Variables of Interest and Network Variability during**

**Resting State + Group Interaction Terms:** Significant predictor variables and group-by-network variability interactions during restidng state, where dots represent individual data points. Statistical significance is denoted by asterisks on each graph (*p < 0.05, **p < 0.01, ***p < 0.001). Group-by-network variability interactions revealed divergent relationships across schizophrenia, autism, and TDC in the subcortical, somatomotor, and cingulo-opercular networks with BSFS total and MCCB composite scores.

**Table S6: Statistical Outputs Assessing the Relationship Between Network-Variability and Variables of Interest + Group Interaction Terms during Resting State**

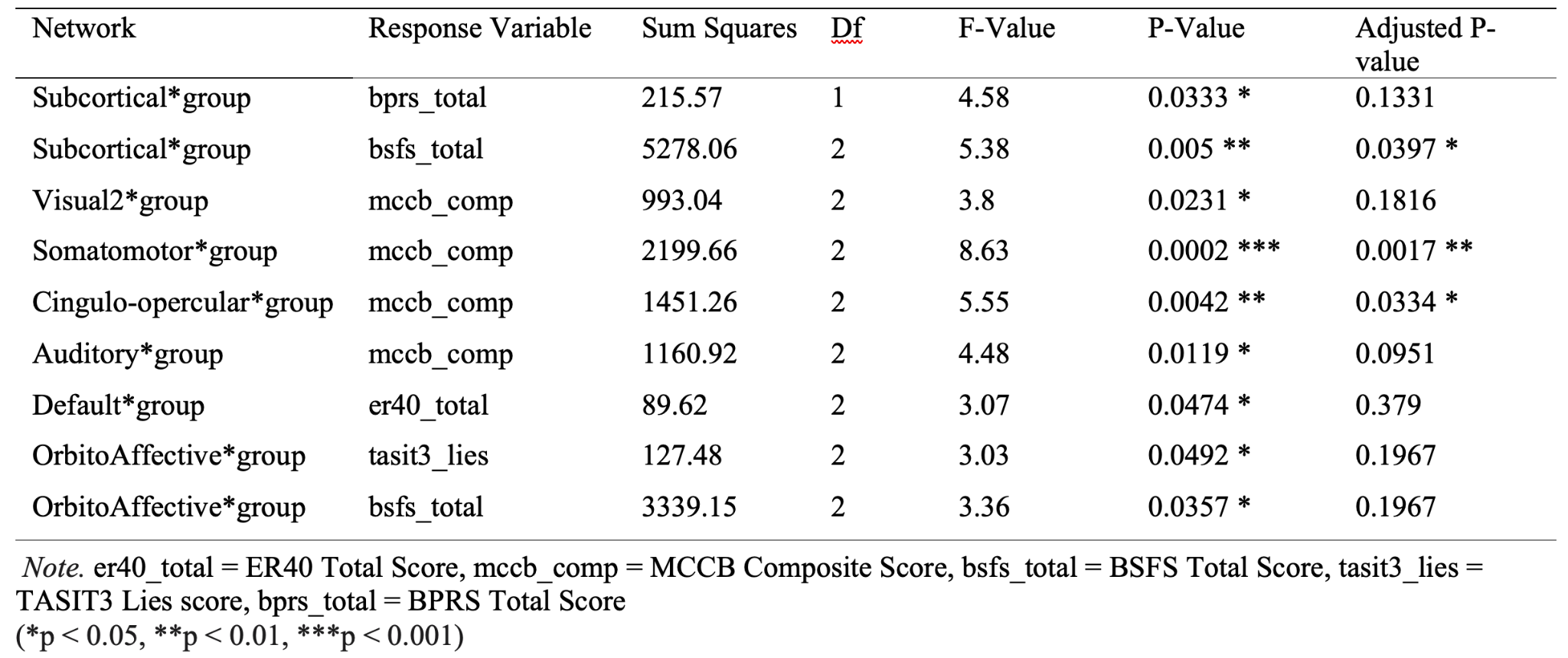

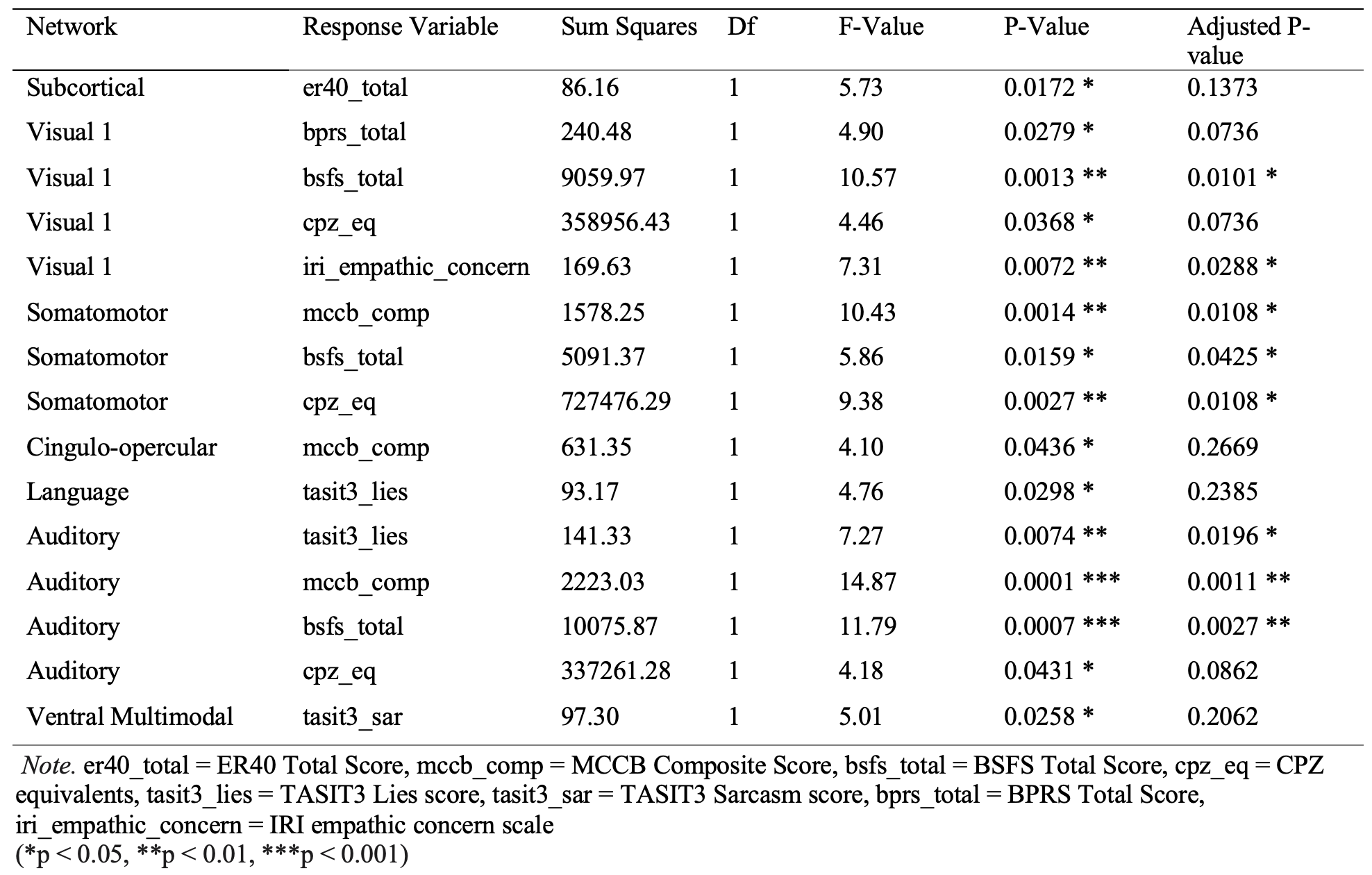
**Table S7: Statistical Outputs Assessing the Relationship Between Network-Variability and Variables of Interest during EA Task**

**References**

1.	First MB, Spitzer RL, Gibbon M, Williams JBW, Others. Structured clinical interview for DSM-IV-TR axis I disorders, research version, patient edition. SCID-I/P New York, NY, USA:; 2002.

2.	Lord C, Rutter M, DiLavore P, Risi S, Gotham K, Bishop S. Autism diagnostic observation schedule–2nd edition (ADOS-2). Los Angeles, CA: Western Psychological Corporation. 2012;284:474–478.

3.	Leucht S, Samara M, Heres S, Davis JM. Dose Equivalents for Antipsychotic Drugs: The DDD Method. Schizophr Bull. 2016;42 Suppl 1:S90–S94.

4.	Wechsler D. Wechsler abbreviated scale of intelligence--second edition. PsycTESTS Dataset. 2018.

5.	Wechsler D. Wechsler Test of Adult Reading: WTAR. Psychological Corporation; 2001.

6.	Kern RS, Penn DL, Lee J, Horan WP, Reise SP, Ochsner KN, et al. Adapting social neuroscience measures for schizophrenia clinical trials, Part 2: trolling the depths of psychometric properties. Schizophr Bull. 2013;39:1201–1210.

7.	Olbert CM, Penn DL, Kern RS, Lee J, Horan WP, Reise SP, et al. Adapting social neuroscience measures for schizophrenia clinical trials, part 3: fathoming external validity. Schizophr Bull. 2013;39:1211–1218.

8.	Esteban O, Markiewicz CJ, Blair RW, Moodie CA, Isik AI, Erramuzpe A, et al. fMRIPrep: a robust preprocessing pipeline for functional MRI. Nat Methods. 2019;16:111–116.

9.	Esteban O, Blair RW, Nielson DM, Varada JC, Marrett S, Thomas AG, et al. Crowdsourced MRI quality metrics and expert quality annotations for training of humans and machines. Sci Data. 2019;6:30.

10.	Avants BB, Epstein CL, Grossman M, Gee JC. Symmetric diffeomorphic image registration with cross-correlation: evaluating automated labeling of elderly and neurodegenerative brain. Med Image Anal. 2008;12:26–41.

11.	Zhang Y, Brady M, Smith S. Segmentation of brain MR images through a hidden Markov random field model and the expectation-maximization algorithm. IEEE Trans Med Imaging. 2001;20:45–57.

12.	Dale AM, Fischl B, Sereno MI. Cortical surface-based analysis. I. Segmentation and surface reconstruction. Neuroimage. 1999;9:179–194.

13.	Treiber JM, White NS, Steed TC, Bartsch H, Holland D, Farid N, et al. Characterization and Correction of Geometric Distortions in 814 Diffusion Weighted Images. PLoS One. 2016;11:e0152472.

14.	Jenkinson M, Bannister P, Brady M, Smith S. Improved optimization for the robust and accurate linear registration and motion correction of brain images. Neuroimage. 2002;17:825–841.

15.	Cox RW, Hyde JS. Software tools for analysis and visualization of fMRI data. NMR Biomed. 1997;10:171–178.

16.	Dickie EW, Anticevic A, Smith DE, Coalson TS, Manogaran M, Calarco N, et al. Ciftify: A framework for surface-based analysis of legacy MR acquisitions. Neuroimage. 2019;197:818–826.

17.	Glasser MF, Sotiropoulos SN, Wilson JA, Coalson TS, Fischl B, Andersson JL, et al. The minimal preprocessing pipelines for the Human Connectome Project. Neuroimage. 2013;80:105–124.

18.	Satterthwaite TD, Elliott MA, Gerraty RT, Ruparel K, Loughead J, Calkins ME, et al. An improved framework for confound regression and filtering for control of motion artifact in the preprocessing of resting-state functional connectivity data. Neuroimage. 2013;64:240–256.

19.	Muschelli J, Nebel MB, Caffo BS, Barber AD, Pekar JJ, Mostofsky SH. Reduction of motion-related artifacts in resting state fMRI using aCompCor. Neuroimage. 2014;96:22–35.
